# Integrative RNA-Seq analysis of *Capsicum annuum L*.*-Phytophthora capsici L.* pathosystem reveals molecular cross-talk and activation of host defence response

**DOI:** 10.1101/2021.04.03.438323

**Authors:** Tilahun Rabuma, Om Prakash Gupta, Manju Yadav, Vinod Chhokar

## Abstract

**Background:** Chili pepper (*Capsicum annuum* L.) being one of an important member of the Solanaceae family, and its productivity is highly affected by the fungal pathogen *Phytophthora capsici* L. Other to CM-344, the unavailability of resistant landraces to all possible strains of *P. capsici* imposes a serious threat to its global production. This is because of our current understanding of the molecular mechanisms associated with the defence response in *C. annuum*-*P. capsici* pathosystem is limited. Therefore, the current study used RNA-seq technology to dissect the genes associated with defence response against *P. capsici* infection in two contrasting landraces, *i.e.* GojamMecha_9086 (Resistant) and Dabat_80045 (susceptible) exposed to *P. capsici* infection.

**Results:** The transcriptomes from 4 leaf samples (RC, RI, SC and SI) of chili pepper resulted in a total of 1,18, 879 assembled transcripts (with a mean TL of 813.23bp and N50 of 1,277bp) along with 52,384 pooled unigenes with (mean UL of 1029.36 bp and N50 of 1,403bp). The enrichment analysis of the transcripts indicated 23 different KEGG pathways under five main categories. Further, 774 and 484 differentially expressed genes (DEGs) were obtained from RC vs. RI and SC vs. SI leaf samples, respectively. Of these, 57 DEGs were found to be associated with defence responses against *P. capsici* infection. The defence-related genes, such as *LTPL*, defensin J1-2-like, peroxidase 5-like, UGT, and GRP proteins-like, were more significantly upregulated in RC vs. RI. Furthermore, RT-qPCR analysis of six randomly selected genes validated the results of Illumina NextSeq500 sequencing results. Furthermore, a total of 58 TF families (bHLH most abundant) and 2,095 protein families (Protein kinase, PF00069, most abundant) were observed across all the samples with maximum hits in RI and SI samples.

**Conclusions:** RNA-Seq analysis of chili pepper’s during *P.capsici* infection revealed differential regulation of genes associated with defence and signaling response with shared coordination of molecular function, cellular component and biological processing. The results presented here would enhance our present understanding of the defence response in chili pepper against *P. capsici* infection, which could be utilized by the molecular breeders to develop resistant chili genotypes.

## Background

Chili pepper (*Capsicum annuum* L.) belongs to the Solanaceae family under the *Capsicum* genus. This genus consists of several important pepper species such as *C. annuum* L., *C. baccatum* L., *C. chinense* L., *C. frutescens* L. and *C. pubescens* L. etc. The chili pepper is native to central and South America and was introduced to the rest part of the world by traders (Pickersgill, 1997). It has a diverse food usage such as spice, consumed fresh, in powder form (Dias *et al*., 2013; Wahyuni *et al*., 2013) and medicinal uses such as drugs, condiments, ointments and relief of pain (Pawar *et al*., 2011; Chamikara *et al*., 2016). Indian states such as Andhra Pradesh, Karnataka, Maharashtra, Orissa and Tamil Nadu account for more than 75% of the area and total production (Reddy *et al*.,2014). In Ethiopia, pepper is cultivated as an income source in several parts of the country and is an important constituent of traditional food because of its pungency and colour (Gebretsadkan *et al*., 2018). Although the economic importance of chili pepper in human nutrition is high, the productivity of this crop is highly affected by both biotic and abiotic factors. Among the biotic factors, fungal pathogens are the most economically important pepper pathogens that trigger diseases like downy mildew, *Phytophthora* late blight, collar rot, and root rot, causing severe yield loss (Yin *et al*., 2012). Among the fungal pathogens, *Phytophthora capsici* L. has been identified as one of the major pathogens limiting its production. *P. capsici* is soil-borne and grouped under the class of oomycete fungi causing the *Phytophthora* stem, collar and root rots and crown blight disease in *Capsicum* species (Barksdale *et al*., 1984; Ristaino and Johnston, 1999; Walker and Bosland, 1999). This pathogen has a broad host range in the Solanaceae (tomato, eggplant, pepper) and Cucurbitaceae (squash, pumpkin, zucchini, cucumber and watermelon) families (Ristaino and Johnston, 1999). Being one of the major pathogen of chili pepper worldwide, current management strategies are not effective (Barchenger *et al*., 2018).

Infection caused by *P. capsici* in chili pepper is distributed to all parts of the plant, including roots, stems, leaves and fruit at any stage of growth (Oelke *et al*. 2003). The CM-334, the most studied resistant landrace of chili pepper, is resistant to root rot, foliar blight, and stem blight (Walker and Bosland, 1999). Owing to different combinations of resistance genes, CM334 is highly resistant to all of the diseases caused even by the most virulent strains of *P. capsici* (Sy *et al*., 2008) and has been widely used in breeding programs (Barbosa and Bosland, 2008). However, to date, no CM-334 derived lines have been developed that could consistently show resistance to all disease symptoms (Oelke *et al*., 2003; Sy *et al*., 2008). The available landraces of resistant pepper other than the CM-334 landrace are not equally resistant to all races of the pathogens and had failed to resist all the infections initiated by the pathogen (Conner *et al*., 2010). Due to physiological, genetic and molecular complexities, investigating the resistance mechanism in *C. annuum* against *P. capsici* is quite challenging (Quirin *et al*., 2005). In addition, improving chili pepper is mainly hindered by the infrequent availability of genetic information and host defence mechanisms underlying disease resistance (Marame *et al.,* 2009). The understanding of the disease response in pepper against *P.capsici* infection is limited (Shen *et al*., 2015). The development of NGS technology coupled with high-precision bioinformatics technologies has facilitated the understanding of the molecular mechanism of foot rot (Hao *et al*., 2016). Shen *et al*. (2015) used RNA-sequencing technology to identify differentially expressed genes associated with defence responses against *P. capsici* in the resistance line PI201234. Liu and co-workers (Liu et al., 2013) demonstrated the capsaicinoids biosynthetic pathway in chili pepper using RNA-Seq approach. Kim and coworkers (Kim et al., 2017) generated a new reference genome of hot pepper and showed the massive evolution of plant disease-resistance genes by retro duplication. Recently, a novel RGA-based marker analysis was performed between two contrasting *Capsicum* germplasms, *i.e.* resistant and susceptible upon infection with *P. capsici* (Jundae *et al*., 2019). Despite the numerous NGS-based studies, the differential gene expression and resistance mechanism associated with defence responses against *P. capsici* infection in resistant and susceptible landraces of chili pepper are significantly limited. In light of this, it is quite essential to perform more robust screening of germplasms using NGS technologies, including transcriptomics. Transcriptome sequencing would generate invaluable information about the mechanism of defence response of *P. capsici* infection in chili pepper.

Therefore, in the present study, we performed RNA-Seq analysis of previously identified resistance (GojamMecha_9086) and susceptible (Dabat_80045) landraces (T. Rabuma *et al*., 2020) exposed to *P. capsici* infection and revealed a variety of differentially expressed genes associated with defence responses.

## Results

### Phenotypic characterization of inoculated chili pepper

Before performing RNA-Seq analysis of resistant and susceptible chili pepper landraces, we carried out the screening of available Indian and Ethiopian landraces for resistance and susceptibility. A total of 233, i.e. 148 Ethiopian and 85 Indian landraces, were used for phenotypic screening upon infection with *P. capsici* (Rabuma*, et al*., 2020). From phenotypic characterization, we selected two contrasting landraces, GojamMecha_9086 (resistant; MDI-1.4) and Dabat_80045 (susceptible; MDI-8.6) for transcriptome analysis based on the disease severity index value and phenotypic performance against pathogen infection. The results indicated no obvious symptoms in the resistant landrace GojamMecha_9086 on leaves, but little lesion was observed on the root (Fig 1A&B). In contrast, the brown lesions were observed on the root, crown root and lower part of the plant along with leaf wilting in susceptible landrace Dabat_80045(Fig 1C & D).

**Fig 1:**
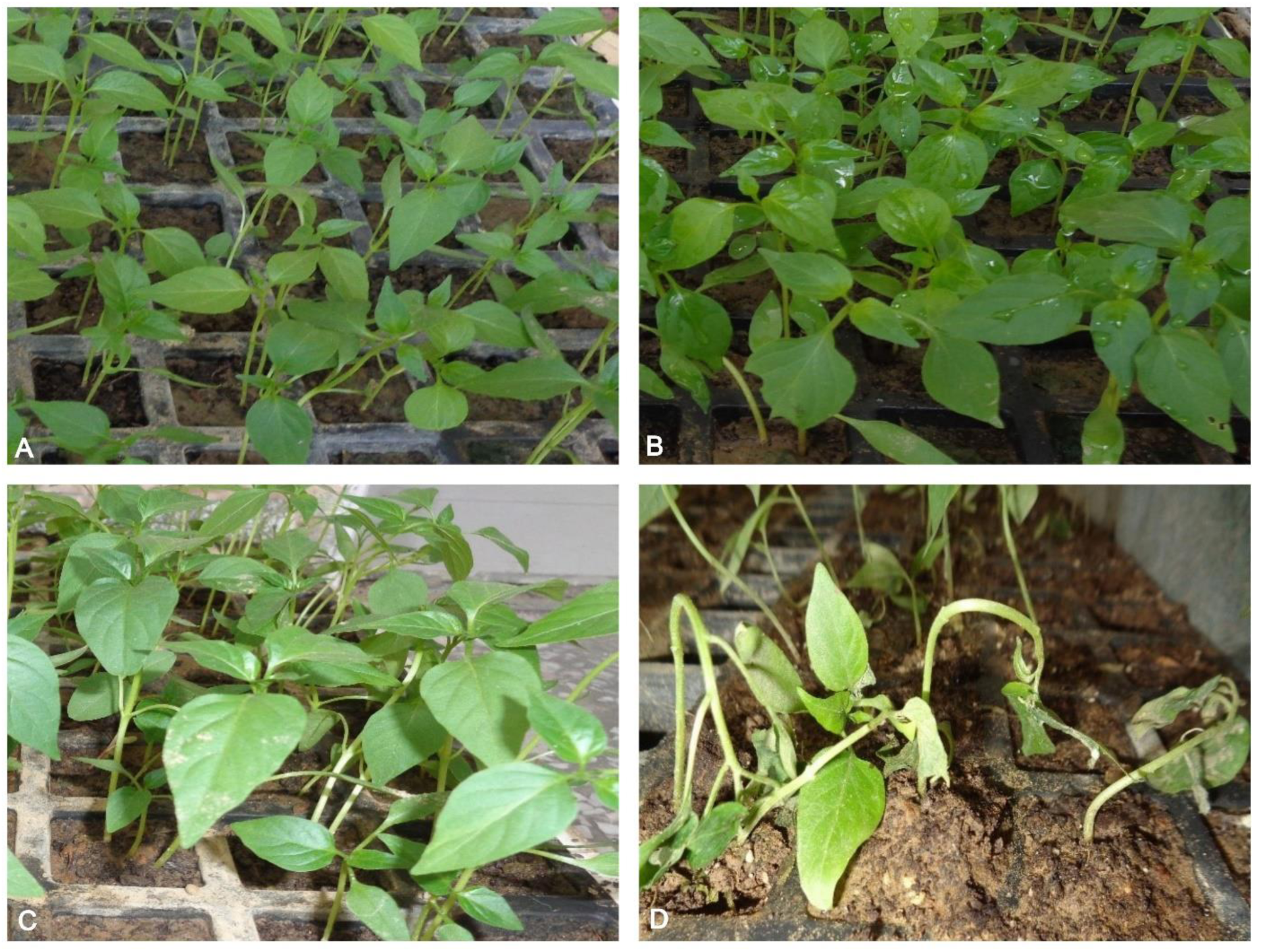
Phenotypic characterization of chili pepper exposed to *P.capsici.* A) GojamMecha_9086 control; B) GojamMecha_9086 infected; C) Dabat_80045 Control; D) Dabat_80045 infected.

### Transcriptome sequence statistics and *De novo* transcriptome assembly

Transcriptome sequencing analysis was carried out to reveal differential expression of defence-related genes in resistant and susceptible chili pepper leaf samples under *P. capsici* infection. A total of 4 libraries (RC, RI, SC, SI) were constructed and sequenced. Illumina NextSeq500 sequencing generated on an average of 22 million high quality (QV>20) paired-end raw reads (∼3.35 GB) from each library and used for downstream analysis after pre-processing. The RNA-Seq raw data of 4 samples were processed to obtained high-quality paired-end reads (Table 1). Pooled assembly from all four libraries was performed, which generated 1,18, 879 assembled transcripts with a mean transcript length of 813.23 and N50 of 1,277 bp (Table 1). The assembled transcripts were further clustered together, resulted in 52,384 pooled unigenes with a total and mean length of 53,922,230 and 1029.36 bp (Table 1). The highest unigenes (16,671) were found to be in the length range of 1,000-2,000bp (Table S1). The CDS prediction analysis from pooled unigenes resulted in 25,552 CDS with a total and mean length of 21,689,736 and 848.84bp (Table 1). Reads mapping back to the pooled CDS resulted in 14,509, 19,250, 10,100, and 22,719 CDS in RC, RI, SC and SI samples, respectively (Table 1). For gene functional analysis, the predicted CDS were searched against the NCBI non-redundant (NR) protein database and resulted in the annotation of 23,738 CDS. Out of 25,552 predicted CDS, 1,814 CDS did not show any significant hits and were marked as novel CDS detected in our RNA-Seq data. The majority of the BLAST hits of the CDS were exhibited against *C. annuum* (17833, 75.13%), followed by *C. chinense* (2803, 11.80%) and *C. baccatum* (1896, 7.98%).

**Table 1.**
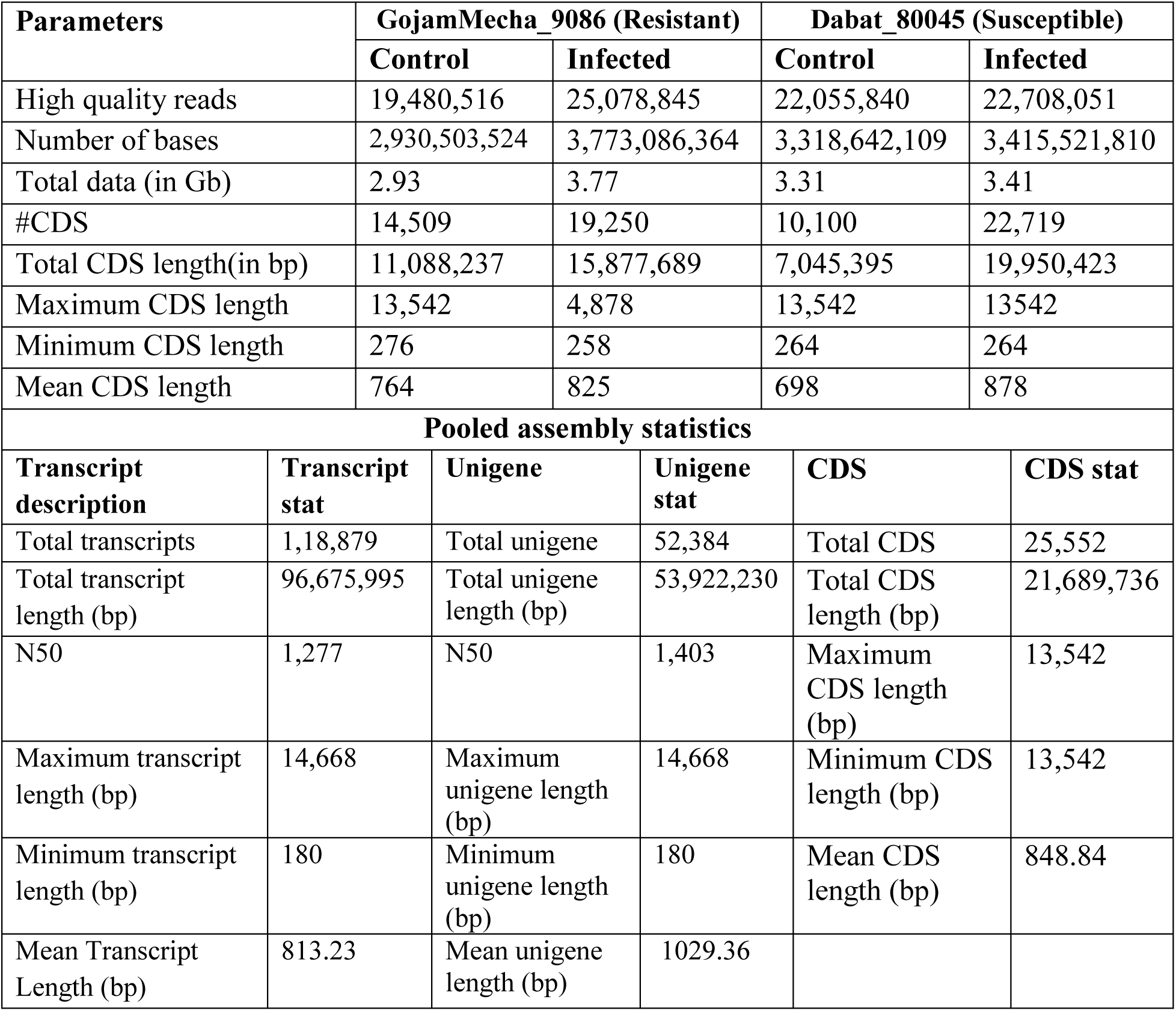
Statistical analysis of RNA-Seq libraries constructed from two contrasting *i.e.* GojamMecha_9086 (resistant) and Dabat_80045 (susceptible) chilli pepper leaf samples exposed to *P. capsici* infection.

### *P. capsici* induced differentially expressed genes (DEGs)

The assembled transcripts were computationally analysed to evaluate differential gene expression in response to *P. capsici* infection. We identified 774 DEGs between RC and RI leaf samples, of which 318 (41.01%) and 456 (58.92%) genes were differentially up and down-regulated (p<0.05), respectively (Table S2). Similarly, 484 DEGs were identified between SC and SI leaf samples, of which 217 (44.84%) and 267 (55.17%) genes were differentially up and down-regulated (p<0.05), respectively (Table S2). Relatively high numbers of up and down-regulated genes were obtained between RC vs. RI leaf than SC vs. SI leaf samples. Among all the DEGs in RC vs. RI, 57 DEGs (45 up and 12 down-regulated) were identified to be associated with the defence response against *P. capsici* infection (Table S3). Similarly, among 484DEGs in SC vs. SI, 29 DEGs were related to defence responses, with 18 and 11 transcripts being down- and up-regulated, respectively. Amongst defence-related genes such as putative late blight resistance protein homolog R1B-13, nonspecific lipid transfer protein GPI-anchored 2-like, L-ascorbate peroxidase 3, peroxisomal-like, ethylene-responsive transcription factor CRF6 isoform X2, and pathogenesis-related leaf protein did not show differential expression between SC and SI leaf. However, most of these genes were differentially expressed between RC and RI leaves. Furthermore, based on the previous reports on defence-related genes, the top 50 differentially expressed genes from both the samples set, *i.e.* RC vs. RI and SC vs. SI, were selected to make heatmap along with hierarchal clustering (Fig 2). For instance, genes encoding UDP-glycosyltransferase 82A1 protein, defensin J1-2-like protein, class I heat shock protein-like, glycine-rich cell wall structural protein-like, and chalcone synthase 1B enzyme were up-regulated. Additionally, Volcano Plot was prepared to depict the graphical representation and distribution of differentially expressed genes between control and treated samples. The ‘volcano plot’ arranges expressed genes along dimensions of biological and statistical significance (Fig 3 A&B).

**Fig 2:**
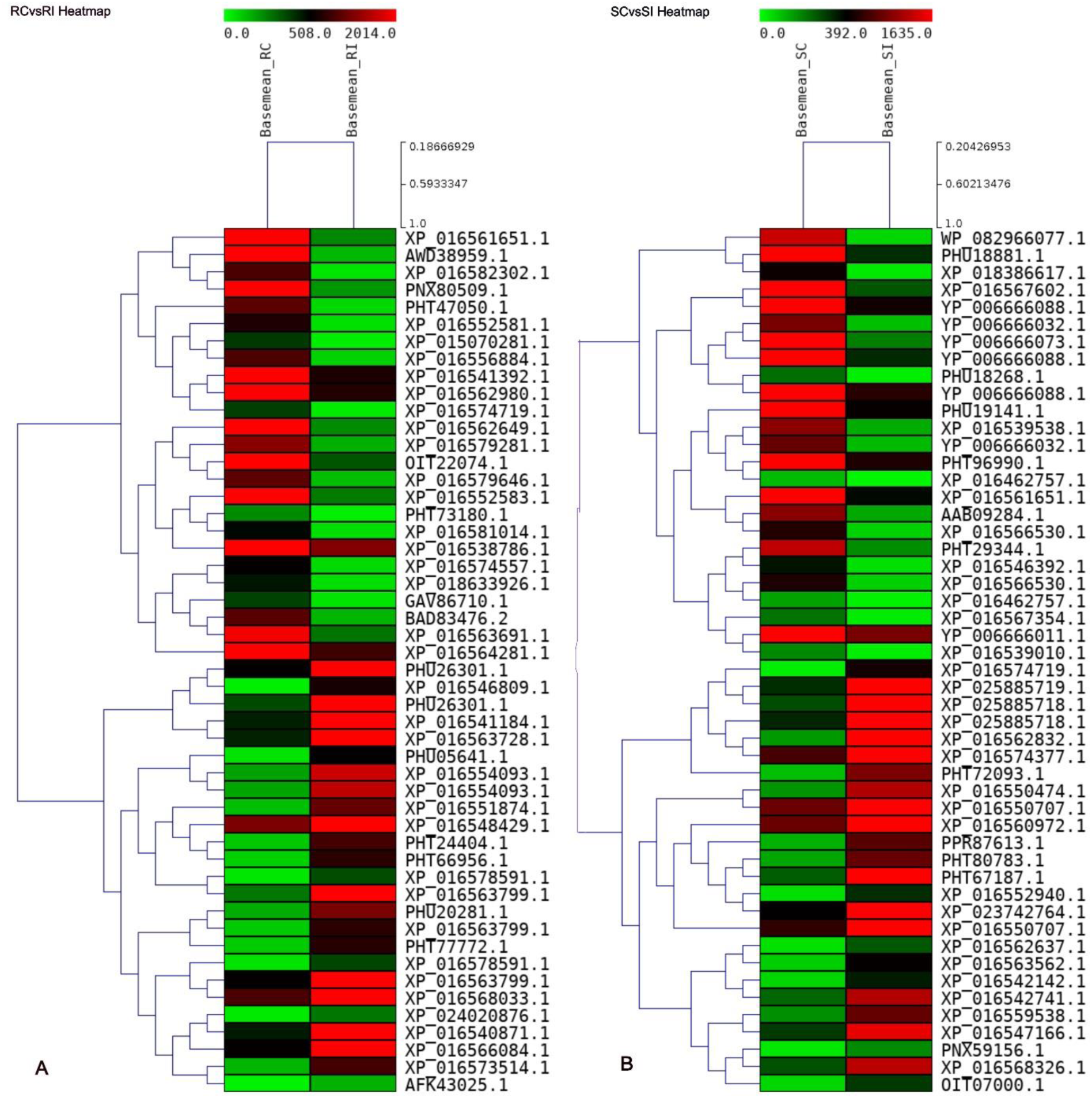
Heat map depicting the top 50 DEGs. The gradient of colours shows, red with highly expressed while green represents no or low expression; A) the normalized gene expression values between RC vs. RI leaf sample, B) the normalized gene expression values between SC vs. SI leaf sample.

**Fig 3:**
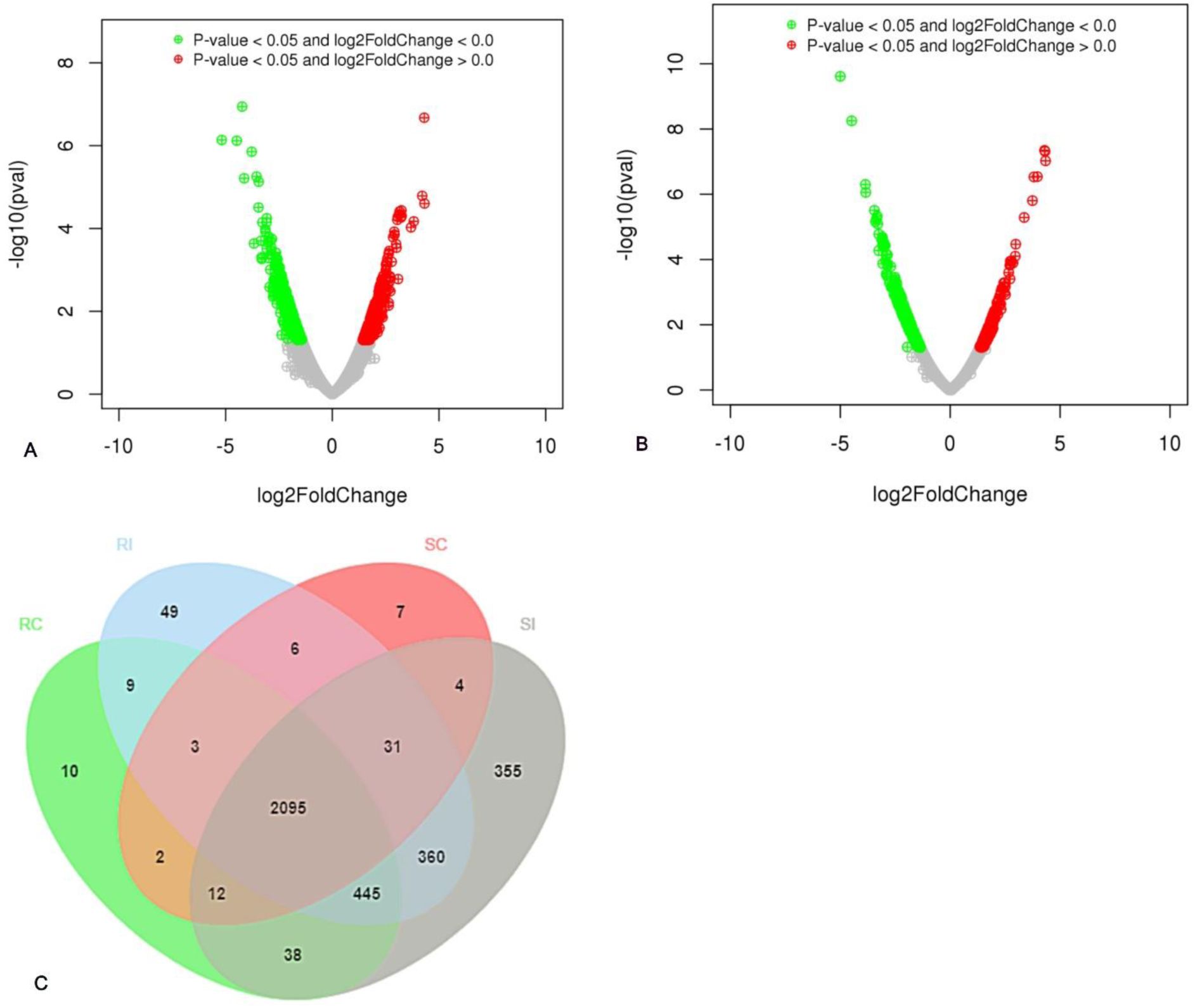
Volcano plot of DEGs between A) RC and RI sample and B) SC and SI sample. The red block on the right side of zero represents the up-regulated genes whereas green block on the left side of zero represents significant down-regulated genes. In addition, the Y-axis represents the negative log of the p-value (<0.05) of the performed statistical test, the data points with low p-values (highly significant) appears towards the top of the plot. In addition, the grey block shows the non-differentially expressed genes; C) Venn diagram showing common DEGs across all the 4 leaf samples, *i.e.* control (RC, SC) and treated (RI, SI).

### GO classification and enrichment analysis of DEGs related to pathosystem

GO mapping was performed to assign the functions to the coding sequence (CDS) predicted in each of the four-leaf samples. BlastX result accession IDs were used to retrieve UniProt IDs making use of the Protein Information Resource (PIR), like PSD, UniProt, SwissProt, TrEMBL, RefSeq, GenPept and PDB databases. BlastX result accession IDs were searched directly in the gene product dbxref table of the GO database. Accordingly, the function of annotated CDS was grouped into three main domains: biological processing (BP), molecular function (MF) and cellular component (CC). A total of 5,605 and 7,913 CDS were found to be involved in biological process, 4,353 and 6,063 in cellular components, 6,751 and 9,475 in molecular function in RC and RI samples, respectively (Fig 4A; Supplementary Fig S1. & Table S4). Besides, 3,980 and 9,231 CDS were found to be involved in biological process, 3,271 and 7,074 CDS in cellular component and 4,721 and 11,084 in molecular function in SC and SI leaf samples, respectively (Fig 4B; Supplementary Fig S2. & Table S4).

**Fig 4:**
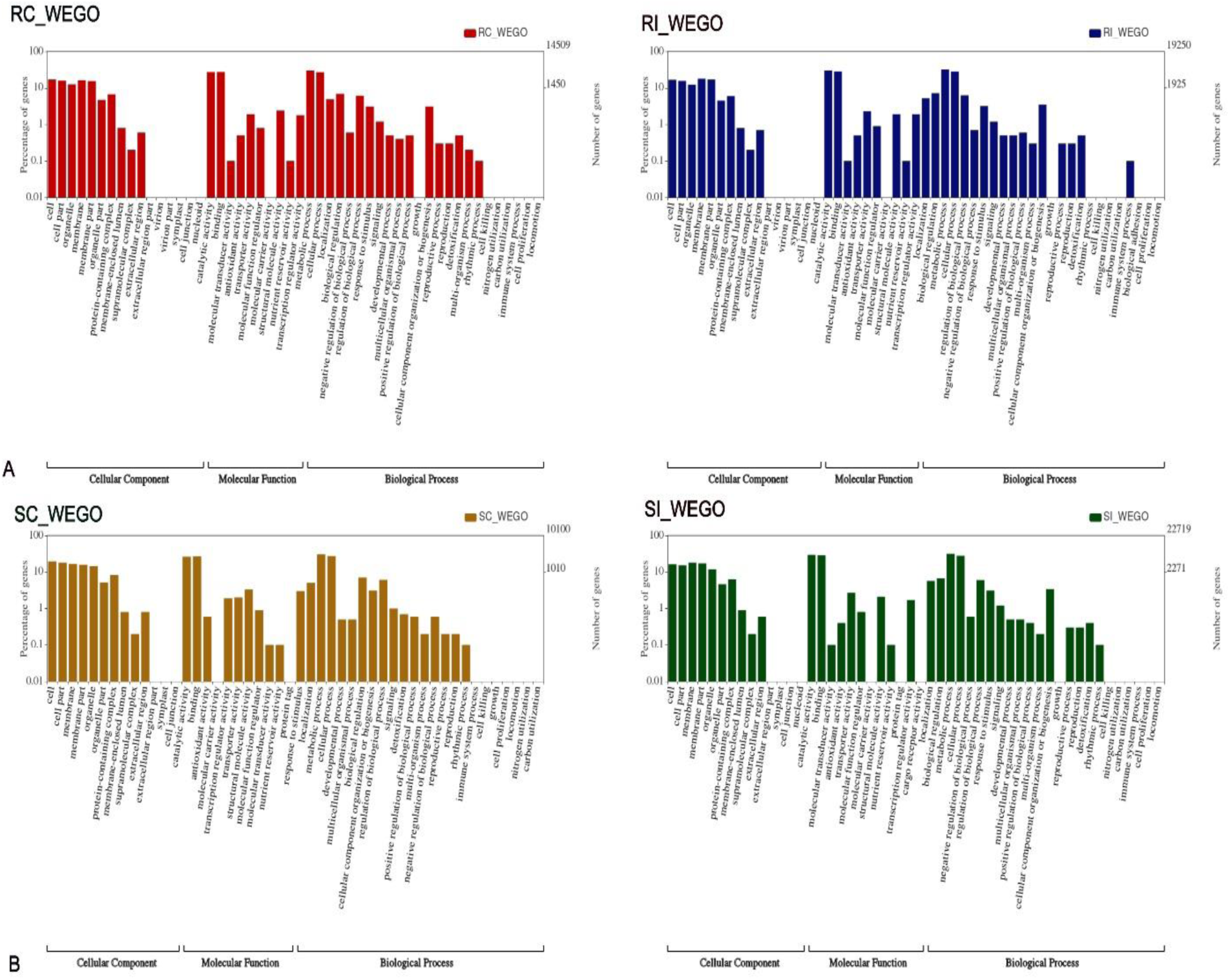
The Web Gene Ontology Annotation Plot (WEGO) showing the distribution of DEGs into three GO tern i.e. cellular, molecular and biological functions. A) RC and RI leaf sample; B) SC and SI leaf samples.

In all the four-leaf samples, the highest number of CDS (112, 159, 74 & 177 in RC, RI, SC & SI, respectively) were shown to be involved in biological processing related to the various organic metabolic process followed by the cellular metabolic process (105, 150, 69,& 171 in RC, RI, SC & SI, respectively). In cellular component, the maximum numbers of CDS were involved in the membrane (2189, 3221, 1552 & 3850 in RC, RI, SC & SI, respectively) followed by an intrinsic component of membrane (1793, 2725, 1381 & 3261 in RC, RI, SC & SI, respectively). Similarly, in molecular function, the maximum number of CDS (993 & 575 in RC & SC respectively) were involved in binding of both organic cyclic compound and heterocyclic compound while 1325 (RI), and 1521 (SI) CDS were involved in transferase activity, which might be related to defence against *P. capsici* pathogens. Annotated genes related to immune system process were detected in only resistance inoculated leaf sample, showing that expression of these genes might be dependent on the inoculation of the pathogens (*P. capsici*) and thus, associated with defence against this pathogen.

### KEGG Pathways mapping, plant transcription factor (pTF) and protein family (Pfam)

To further investigate the mechanism of defence response in chili pepper upon infection with *P. capsici*, KEGG pathway analysis of DEGs was performed in all the four assemblies. The identified CDS were mapped to reference canonical pathways in the KEGG to elucidate the involvement of CDS in various KEGG pathways. The enrichment analysis of the transcripts (4183, 5089, 3320 & 5515 CDS in RC, RI, SC and SI leaf sample, respectively) indicated 23 different KEGG pathways under five main categories (Fig. 5). The SI leaf sample recorded the highest number of genes (5515 CDS) in across five categories of biological pathways and followed by RI sample (5089 CDS) (Fig. 5). The KEGG analysis included KEGG Orthology (KO), enzyme commission (EC) numbers, and metabolic pathways of predicted CDS using KAAS server. Amongst all the pathway identified, a total of 1733, 2197, 1354 and 2324 CDS in RC, RI, SC and SI, respectively, were functionally assigned for metabolism pathway making it the largest KEGG classification group including carbohydrate metabolism, energy metabolism, lipid metabolism, nucleotide metabolism, amino acid metabolism etc. The detailed description including number of CDS in each category and samples is mentioned in (Table S5). Nevertheless, we also predicted TF family to dissect its abundance in all the four assemblies against 320,370 known TFs in the PlantTFDB 4.0 database (http://planttfdb.cbi.pku.edu.cn/) based on E-value<1e-05 via BlastX. The analysis resulted in the identification of a large number of transcription factors families across all the four assemblies (Fig. 6). A total of 58 families, including bHLH, MYB_related, C3H, WRKY, etc., were identified with basic helix loop helix (bHLH) having the highest hits of 714, 1022, 498 & 1222 in RC, RI, SC & SI leaf samples, respectively followed by NAC having 487, 721, 364 & 826 CDS in RC, RI, SC & SI leaf samples, respectively (Table S6). Additionally, protein family (Pfam) prediction was performed to know the abundance of protein families for each CDS in all the 4 assemblies based on E-value<1e-05 via BlastX to Pfam 32.0 databases. Results indicated an average of ∼3,982 families with ∼2095 common protein families across all the four assemblies of chili pepper upon *P. capsici* (Fig. 7). The distribution of top ten predicted protein families revealed that protein kinase domain (PF00069) was the most abundant family (290, 489, 160 & 571 CDS in RC, RI, SC & SI, respectively followed by RNA binding protein (PF00076) domain (184, 224, 130, & 255 in RC, RI, SC & SI, respectively) and protein tyrosine kinase (PF07714) (Fig. 7).

**Fig 5:**
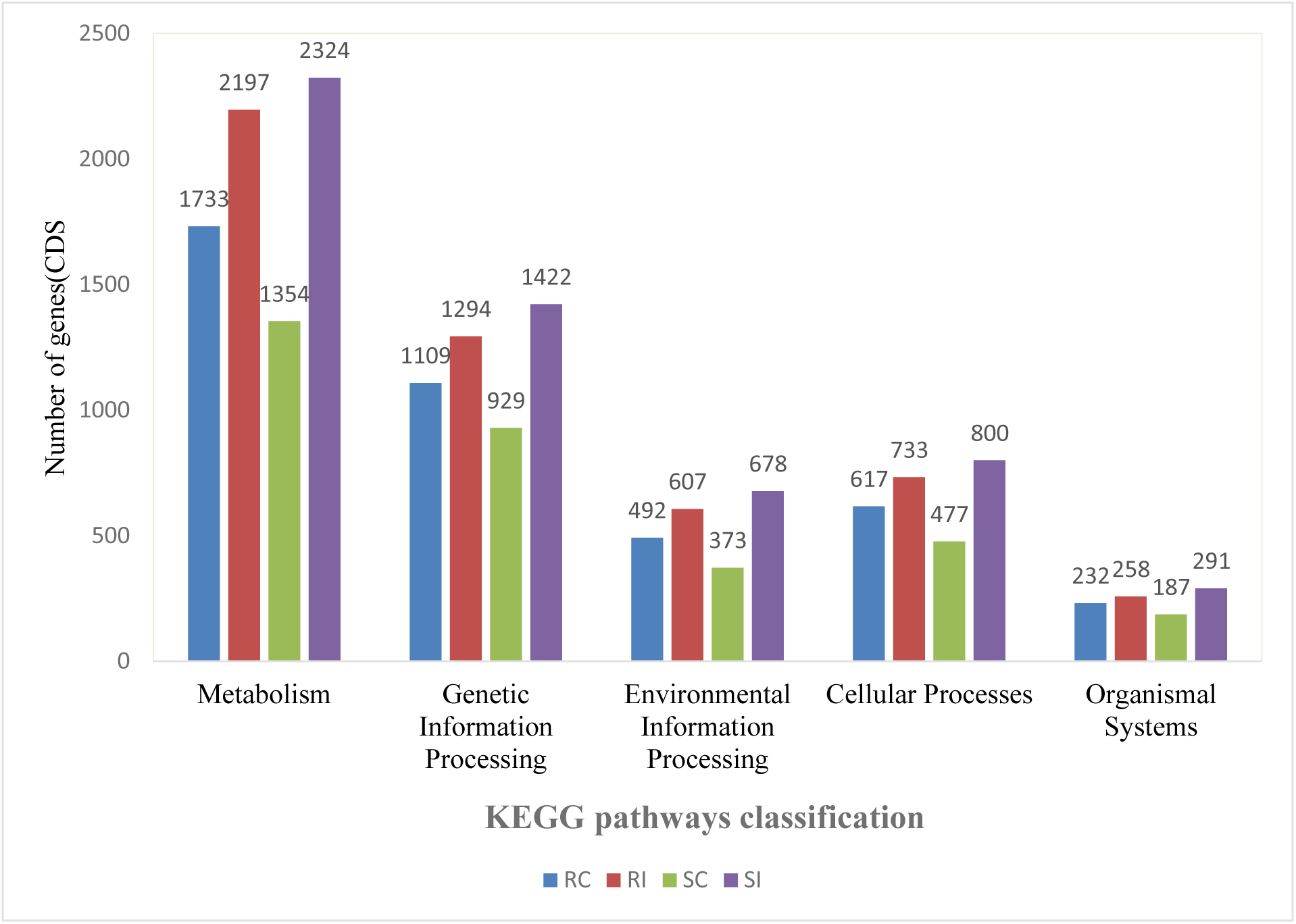
Distribution of top 5 highly enriched KEGG pathway in resistant and susceptible landraces of chili peppers exposed to *P. capsici* infection. The vertical y-axis represents the number of CDS involved in each pathway and the x-axis represents the name of the pathways.

**Fig 6:**
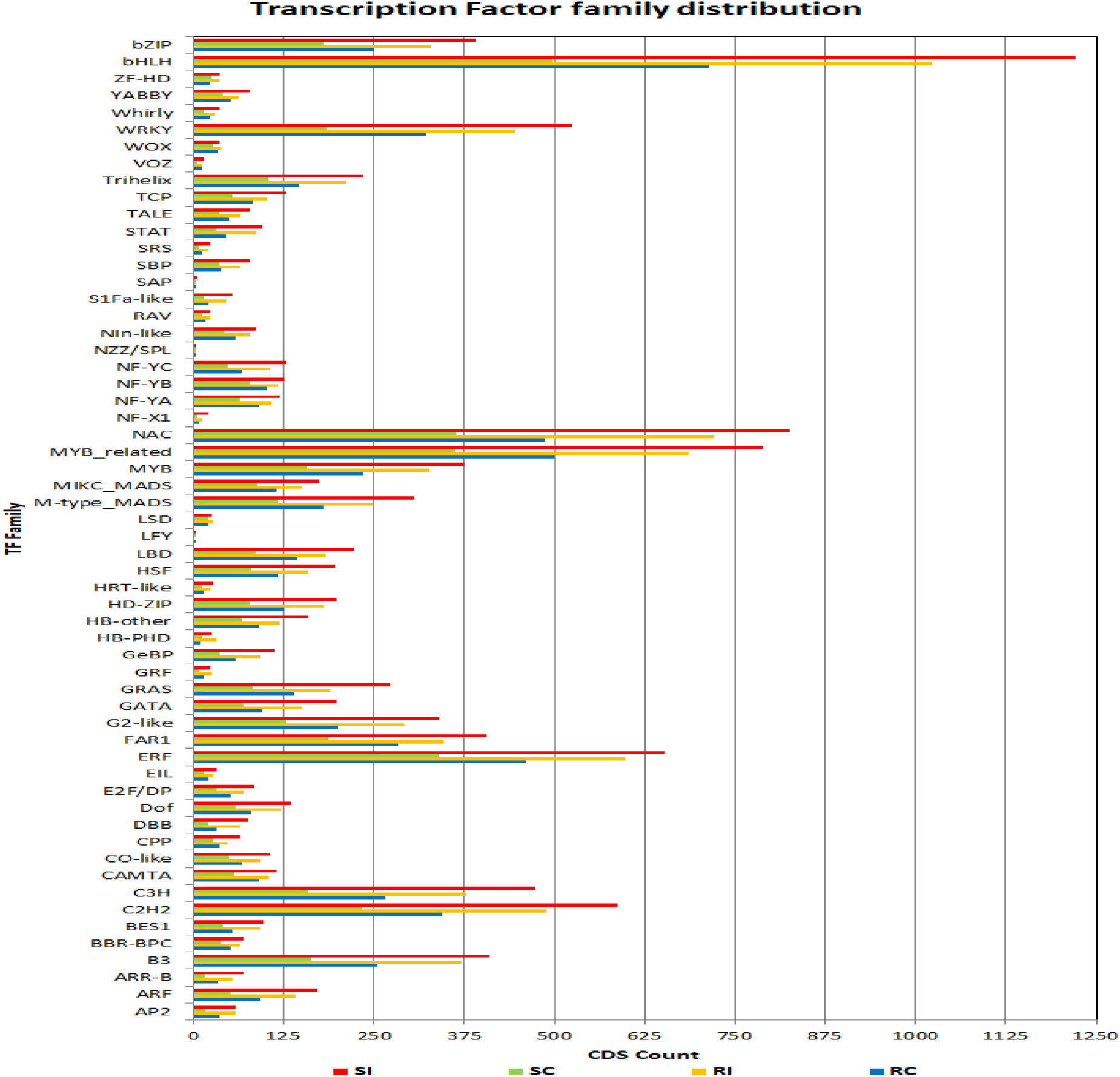
Distribution of significantly enriched transcription factors families in resistant and susceptible landraces of chili peppers exposed to *P. capsici* infection.

**Fig 7:**
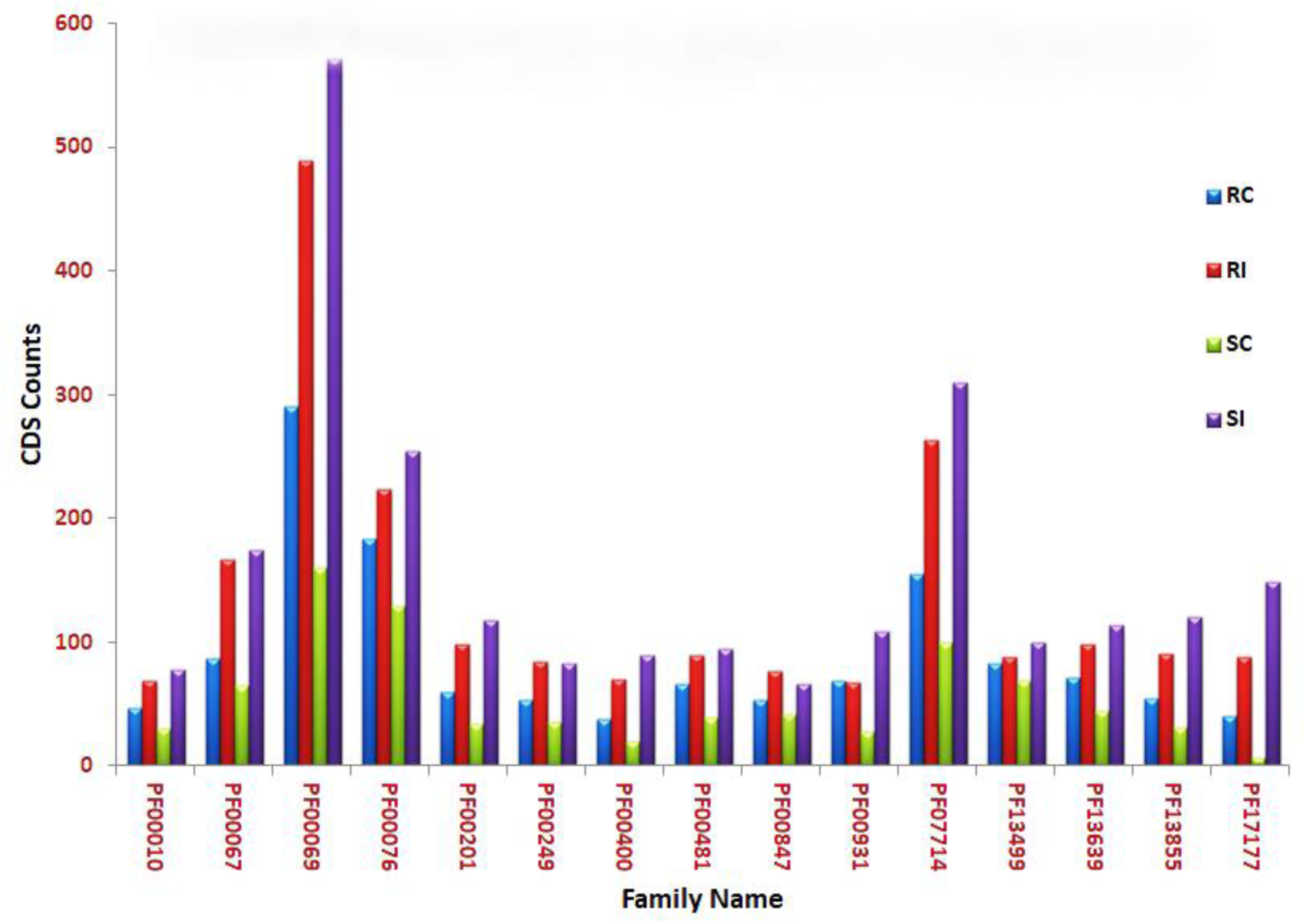
Distribution of the top 15 most abundant protein families in resistant and susceptible landraces of chili peppers exposed to *P. capsici* infection.

### Mining of SSRs markers associated *P.capsici* infection

SSRs, also known as microsatellites, are tandem repeat motifs of 1–6 bases and serve as the key molecular markers in population and conservation genetics, molecular epidemiology and pathology, and gene mapping. In the present study, SSRs were detected using MIcroSAtellite Identification Tool (MISA v1.0) from the pooled set of unigenes. The potential SSRs were identified as ranging from dinucleotide motifs with a minimum of ten repeats, trinucleotide motif with a minimum of four repeats, tetra, penta and hexanucleotide motifs with a minimum of five repeats. A maximum distance of 200 nucleotides was allowed between two SSRs leading to 7,233 SSR from 52,384 pooled unigene sequences. Based on upstream and downstream 200 bp flanking sequence, the SSRs were validated *in silico* and finally, 3,174 SSRs were obtained (Table S7). The analysis result depicted that tri-nucleotide repeats were the most abundant (6,755) SSRs in all the four-leaf samples followed by di-nucleotide repeats with 288 counts. Similarly, motif prediction of these SSR showed an abundance of T/A followed by CT/GA in all four assemblies.

### Validation of expression of selected DEGs

Transcriptome sequencing result obtained from Illumina NextSeq500 PE was validated using RT-qPCR by randomly selecting six genes from the same RNA used for RNASeq analysis. The qRT-PCR data were found consistent with the DEG output in both resistant and susceptible genotype. Among the RT-qPCR analysed genes, all the six transcripts exhibited the same expression pattern with transcriptomic DEG output (Fig. 8 A-F). In resistant genotype GojamMecha_9086 upon infection, all of the six transcripts, i.e. PLTP(FC: 1.96, P-value: 0.02), EP3(FC:2.08, P-value:0.02), Defensin J1-2 like(FC:3.69, P-value:0.00), PRP(FC:0.78, P-value:0.04), LTLP(FC:3.12, P-value:0.00), and ERTF(FC:1.77, P-value:0.00) genes were differentially up-regulated while in susceptible genotype Dabat_80045, PRP(FC:1.22, P-value:0.05), and LTLP(FC:0.38, P-value:0.04) genes were up regulated whereas PLTP(FC:-1.66, P-value:0.03) and Defensin J1-2 like(FC:-0.70, P-value:0.02) were down regulated. The RT-qPCR amplified product of five differentially expressed genes from control and treated samples are shown in Fig S3(Fig S3 Original Gel image).

**Fig 8:**
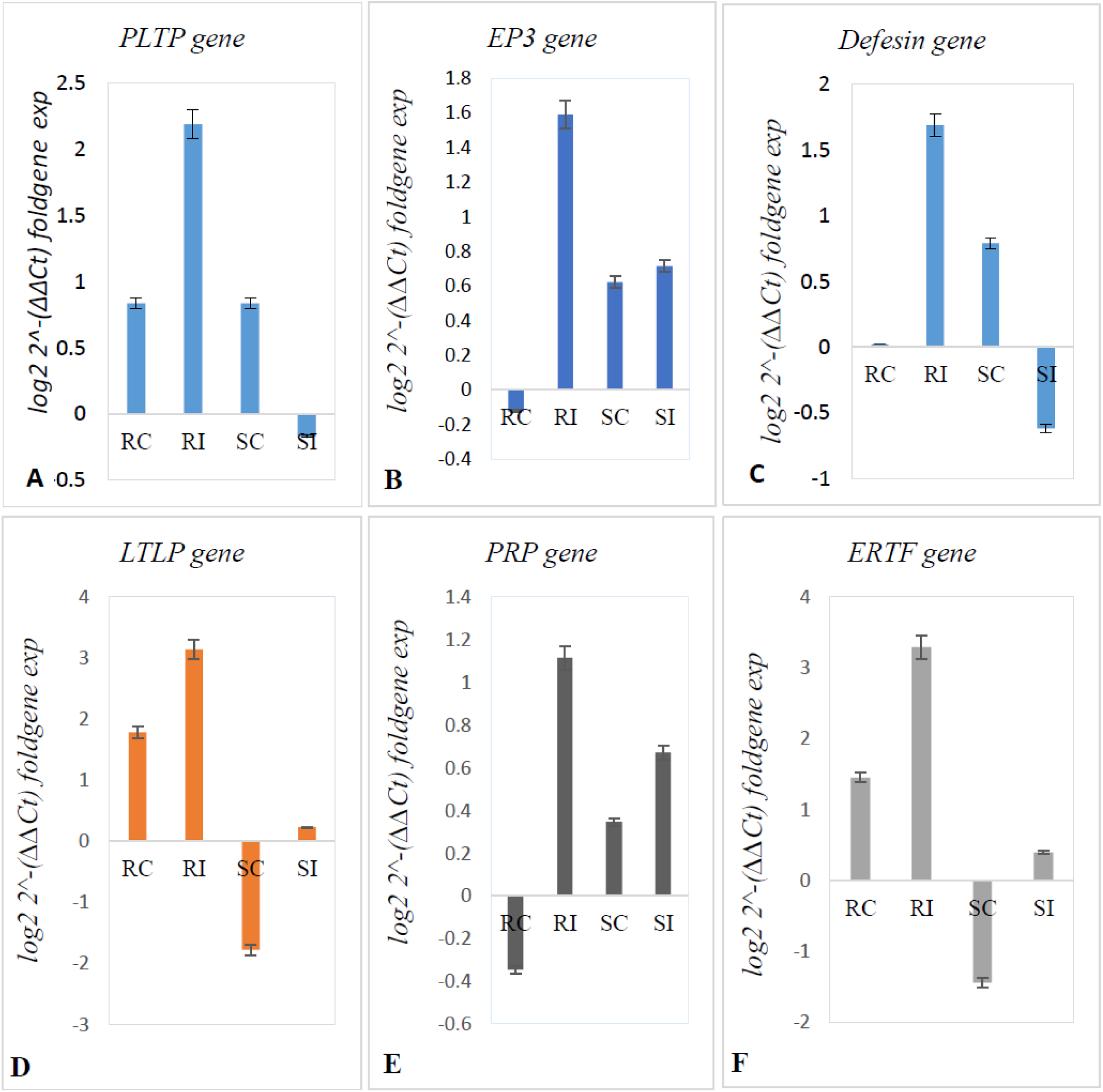
qRT-PCR validation of six key DEGs associated with defence response upon *P. capsici* infection. A) putative lipid transfer protein DIR1 gene (PLTP), B) endochitinase EP3-like gene(EP-3), C) defensin J1-2-like gene (Defensin), D) lipid transfer-like protein VAS gene(LTLP), E) pathogenesis-related protein STH-2-like gene(PRP), and F) ethylene-responsive transcription factor RAP2-7-like isoform X2 gene (ERTF). The y-axis represents the log fold relative gene expression and the x-axis represents the chili pepper leaf samples (RC, RI, SC, and SI). Actin-7 like gene was used as an internal control gene for data normalization. The experiment was replicated three times and the resulting data was presented with errors bar, with n=3.

## Discussion

Despite one of the commercial spices, the productivity of *C. annuum* L. is highly affected by several pathogens including *P. capsici.* The understanding of the disease resistance gene in *C. annuum* in response to *P. capsici* infection is limited (Shen *et al*., 2015). In the recent years, RNA-Seq approach has emerged as one of the key NGS techniques for rapid and efficient characterization of differential gene expression associated with pathogen infection under control and treated condition (Wang *et al*., 2010). It provides a valuable resource for the elucidation of molecular mechanisms and candidate genes for functional analysis (Wang *et al*., 2019). The present study was designed to carry out the RNA-Seq analysis of two contrasting (resistant and susceptible) chili pepper landrace upon infection with *P. capsici* to decipher the differentially expressed genes associated with defence response.

RNA-Seq data resulted in 1,18,879 assembled transcripts with mean transcript length of 813.23 and N50 of 1,277 bp, which is higher than 47,575 assembled transcripts with an average length of 1437.22 bp reported by Shen *et al*. (2015) in pepper line PI 201234. Similarly, we generated ∼52,384-pooled unigenes from four libraries compared to 116,432 unigenes obtained from six libraries of *P. nigrum* L. in response to *P. capsici* infection (Hao *et al*., 2016). About 92.9 and 75.13% of the CDS were functionally annotated and matching with available *C. annuum* reference genome, respectively, representing a high degree coverage and functionality of our RNA-Seq data.

The Gene Ontology mapping showed that metabolic processing, membrane, and binding terms were the dominant in biological process, cellular component and molecular function, respectively, which was not in agreement with the report in Hao *et al*. (2016) which might be due to different genetic background of the selected genotypes for resistance and susceptibility level to *P. capsici*. Predicted CDS genes assigned to molecular function were more enriched in RI (9,475 CDS) and SI (11,084 CDS) samples relative to their corresponding control signifying their role during infection which was agreed with the Richins *et al*. (2010). This suggests that genes of catalase activity in resistance genotype (CM334) and glutathione S-transferase in susceptible genotype (NM6-4) were associated with defence response against *P. capsici* infection. In all four-leaf samples, a large number of genes including pepper 9-lipoxygenase involved organic metabolic process were reported which *is* crucial for lipid peroxidation processes during plant defence responses to pathogen infection (Hwang and Hwang, 2010). Reports have suggested the active role of secondary metabolites such as phenylpropanoid, and terpenoids during a counter-attack against invading *P. capsici* in black pepper (*Piper nigrum* L.) (Hao *et al*., 2016). Metabolic pathway genes were activated during pathogen infection and abiotic stress, as in the plant senescence process (Pageau *et al*., 2006). Carbohydrate metabolism in the metabolic pathway is the largest KEGG pathway group observed in the present result, which signifies their active role during defence response against infection, which is lying with the work of Zhou *et al*., (2018) where they assigned a maximum number of unigenes to carbohydrate metabolism in *G. littoralis*. Also, our result is supported by the Shen et al., (2015) signifying that higher accumulation of CDS related to terpenoids, polyketides metabolism, secondary metabolites, and xenobiotics biodegradation and metabolism in *P.capsici* treated samples compared to control might be playing an important during interaction with the pathogen.

The RNA seq obtained 152 genes directly associated with resistance against *P. capsici* infection. Among these, lignin-forming anionic peroxidase-like precursor, ethylene-responsive transcription factor RAP2-7-like isoform, putative lipid-transfer protein DIR1, lipid transfer-like protein VAS, Acidic endochitinase Q, endochitinase EP3-like, defensin J1-2-like, UDP-glycosyltransferase 82A1, glycine-rich cell wall structural protein-like, phenylalanine ammonia-lyase, non-specific lipid-transfer protein-like protein At5g64080 isoform X2, non-specific lipid transfer protein GPI-anchored 2-like, lipid transfer protein EARLI 1-like and GDSL esterase/lipase APG-like were differentially up-regulated under *P. capsici* infection in resistant genotype. However, in susceptible genotype, the lipid-transfer protein gene families such as non-specific lipid-transfer protein-like protein At5g64080 isoform X2, non-specific lipid transfer protein GPI-anchored 2-like, putative lipid-transfer protein DIR1 and non-specific lipid-transfer protein 1 were differentially down-regulated. These pathogenesis-related (PR) genes were jointly involved in resisting against *the P. capsici* infection in the leaf of pepper. Varieties of PR proteins were coordinated by cross-talk stress signals and endogenous plant hormones such as salicylic acid (SA), methyl jasmonate (MeJA), and ethylene (ET) (Kim *et al*., 2019). The effect of many PR genes (polygenes) associated with resistance response against *P. capsici* and the regulatory mechanisms of pepper against the *P. capsici* infection were complex (Shen *et al*., 2015). The plant responds to the pathogens by increasing expression of the number of genes, some of which might be associated with disease resistance (Chang *et al*., 2004). The plant defence response consists of two major pathways including pathogen-associated molecular pattern (PAMP-triggered immunity (PTI)) and effector-triggered immunity (ETI) (Bali *et al*., 2019). Higher expression levels (1,953 DEGs) of transcripts were reported in *P. flaviflorum* (resistance) than *P. nigrum* (susceptible) in response to *P. capsici* (Hao *et al*., 2016). Moreover, overexpression of PR genes like chitinase, glucanase, thaumatin, defensin and theonin have increased the level of defence response in plants against a range of pathogens (Grover *et al*., 2018). Ethylene biosynthesis was distinctly accumulated together with basic pathogenesis-related (CABPR1) mRNA in pepper leaves upon infection with *P. capsici* (Young and Byung, 2000).

Pathogen invasion triggers activation of a series of genes associated with various signalling pathways as a counter-attack vis-à-vis defence mechanism to cope up with the infection. This type of differential cross-talk between pathogen and host explains as why one genotype is resistant while other as susceptible to the same pathogen. Reports have indicated differential regulation of numerous set of genes associated with defence response, including PR genes, hormone homeostasis in peppers against *P. capsici* infection (Shen *et al*., 2015; Kim *et al*., 2019; Zhen-Hui Gong *et al*., 2013). The CanPOD gene was significantly induced in leaves of pepper by *P. capsici* infection (Wang *et al*., 2013a). The CALTPI and CALTPIII genes were predominantly expressed in various pepper tissues infected by *P. capsici* (Jung *et al*., 2003). Lipid transfer protein (LTP) isolated from the seeds of *C. annuum* is known as Ca-LTP1 involved in morphological deformation and changes in the cells of pathogens (Majid *et al*., 2016). It usually resides in the dense vesicles and displays antifungal activity (Fernando *et al*., 2014). We also recorded up-regulation of lipid transfer-like protein VAS (∼3.12 fold) in RC vs. RI, which might be involved in modulating fungal cell wall more efficiently in resistant landrace than susceptible one. Defensins that belong to antimicrobial peptide and overexpression of defensin enhances resistance to fruit specific anthracnose fungus in pepper (Seo *et al*., 2014). Expression of the defensin J1-1 gene was reported during the ripening of fruit and pathogen wounded tissue (Seo *et al*., 2014). Most plant defensins are involved in defence against the fungal pathogen (Wong *et al*., 2007). Most identified crop plant defensins have been overexpressed under pathogen attack, wound and some abiotic stress leading to resistance (Beer and Vivier, 2011). Defensin inhibits fungal growth by initial binding of the fungal membrane due to electrostatic and/or hydrophobic interactions (Seo *et al*., 2014). Our result showed increased expression (∼3.69 fold) of defensins J1-2 like protein in resistant landrace compared to susceptible one showing its role in resistance *P. capsicum* treated genotype.

Ethylene response factor (ERF) is a multi-member family, which have been reported for their key role during plant-pathogen interaction. One member, CaPTI1 is shown to be up-regulated and involved during defence response in pepper against *P. capsici* (Jin *et al*., 2016). The role of ERF in defence during *P. capsici* infection has also been validated by virus-induced gene silencing (VIGS) of CaPTI1, which significantly reduced the expression of CaPR1, CaDEF1 and CaSAR82 (Jin *et al*., 2016). Our result showed increased expression of ERF gene RAP2-7-like (∼1.77 fold) upon infection in resistant landrace with *P. capsici* supporting its probable role during modulating the defence response in resistant landrace. Similarly, UDP-glycosyltransferase is shown to be associated with modulating the tolerance of plants against various biotic and abiotic stresses (Zhang *et al*., 2014). Xie and co-workers (2016) have documented the induced accumulation of the gene encoding UDP-glycosyltransferase in (*C. annuum)* exposed to 24-epibrassinolide (EBR). UDP-glycosyltransferase 82A1 was found to be up-regulated (∼3.82 fold) in resistant landrace compared to susceptible one suggesting active metabolite flux channelling towards the development of resistance phenotype.

Different transcription factors such as MYB, bHLH and NAC are involved in modulating the gene cascade during various developmental stages and defence response against biotic and abiotic stresses (Agarwal *et al*., 2006; Zhang *et al*., 2009; Kikuchi *et al*., 2013; Deeba *et al*., 2016). Our RNA-Seq data enabled us to identify a total of 58 TFs amongst which bHLH, MYB and NAC were the most abundant ones having differential expression in both resistant and susceptible landraces which might be involved in regulating the defence-related signalling pathway upon *P. capsici* infection.

## Conclusion

In this study, Illumina NextSeq500 of RC, RI, SC, and SI leaf samples generated ∼22 million high quality reads. From the transcriptome sequencing analysis, the genes associated with defence response to *P. capsici* infection were identified in the leaf sample of *C. annuum*. Among the total differentially expressed genes expressed under *C. annuum-P. capsici* pathosystem, 57 DEGs were found to be associated with defence response against *P. capsici* infection between RC and RI leaf samples. It was shown that more number of genes involved in defence response against *P. capsici* were differentially expressed in RI leaf sample relative to the SI leaf sample. The level of resistance and susceptibility among the chili pepper landraces is largely due to the difference in the level of gene expression and molecular variations in the resistance mechanism. The PR genes, TFs and major protein families were identified in chili pepper leaf under *P. capsici* infections. The current findings could be used as invaluable information about the differentially expressed genes associated with defence response against *P. capsici* infection in chili pepper, helping breeders for selection of landrace containing resistance gene and also could help for transgenic development of *C. annuum* accessions having agronomically important traits like disease resistance and high yield.

## Methods

### Plant growth, fungus inoculation, and RNA isolation

To decipher the defence mechanism operating in chili pepper, we selected two contrasting landrace GojamMecha_9086 (resistant) and Dabat_80045 (susceptible). The previously identified, collected and conserved chili pepper seeds of GojamMecha_9086 with accession number: ETH-9086 and Dabat_80045 with accession number: ETH-80045 were procured from Ethiopian Biodiversity Institute (http://www.ebi.gov.et). The level of resistance and susceptibility of the selected genotypes has already been established (Rabuma *et al*., 2020). The chili plant seedlings were grown in pots containing sterilized soil, sand, and compost with a ratio of 1:1:1 (Andrés *et al*., 2005), at a temperature range of 20-35 °C with a relative humidity of 50-75 % from February till late April 2019 (Rabuma et al., 2020). For transcriptome sequencing, the seedlings of both resistant (GojamMecha_9086 with control & treated) and susceptible (Dabat_80045 with control & treated) genotypes were grown in three biological replications. The resistance landrace GojamMecha_9086 denoted as; RC, resistance control and RI, resistance inoculated and the susceptible landrace Dabat_80045 denoted as; SC, susceptible control and SI, susceptible inoculated were used throughout this research paper. Sporulation followed by inoculation was performed by irrigating the pure suspension culture of *P. capsici* (with isolate ID: NCFT-9415.18) with a concentration of 2 x 10^4^ zoospores/mL (Andrés *et al.,* 2005; Ortega *et al*., 1995) into one-month-old chili pepper seedling leaves and stems (Bosland and Lindsey, 1991). After 5 days post-inoculation (dpi), the 4^th^ and 5^th^ leaves of control and treated leaf samples were collected for RNA isolation as described by Shen *et al*. (2015). The leaf samples from all the three biological replications were harvested separately followed by washing and freezing in liquid nitrogen and kept at −80 °C till further use. Leaf samples constitute four sets; resistant inoculated (RI), susceptible inoculated (SI), resistant control (RC) and susceptible control (SC). The total RNA was isolated using *Quick*-RNA Plant Miniprep Kit (ZYMO Research) as per the manufacturer protocol. The RNA samples from each biological replication were pooled to make 4 libraries (RC, RI, SC and SI) to get maximum representation. The quality and quantity of isolated RNA were checked by 1% agarose gel electrophoresis and Nanodrop spectrophotometer, respectively.

### Illumina NextSeq500 PE library preparation

The RNA-Seq paired-end sequencing libraries were prepared from the QC passed RNA samples using Illumina TruSeq Stranded mRNA Sample Prep kit according to the manufacturer manual. The Poly-A containing mRNA molecules were purified using Poly-T oligo attached magnetic beads. Following purification, the mRNA was fragmented into small pieces using divalent cations under elevated temperature. The first-strand cDNA was synthesised using SuperScript and Act-D mix to facilitate RNA dependent synthesis. The 1^st^ strand cDNA was then synthesized to the second strand using a second strand mix. The dscDNA was then purified using AMPure XP beads followed by A-tailing, adapter ligation and libraries are amplified by PCR with 18 *cycles* to increase the quantity of the library and enrich for properly ligated template strands (Bronner *et al*., 2009). The PCR enriched libraries were analysed on 4200 Tape Station system (Agilent Technologies) using high sensitivity D1000 screen tape as per manufacturer instructions. Finally, for each RC, RI, SC and SI leaf sample, the PE libraries were prepared using TruSeq® Stranded mRNA Library Prep kit as per the manufacturer’s instruction. Bioinformatics data analysis workflow steps undertaken during transcriptome sequencing of *C. annuum* L was summarized in (Fig S4).

### Cluster Generation, RNA-Sequencing and Data filtration

After obtaining the Qubit concentration for the libraries and the mean peak sizes from Agilent Tape Station profile, the PE Illumina libraries were loaded onto NextSeq500 for cluster generation and sequencing. The RNA-sequencing using NextSeq500 was performed using 2×75 bp chemistry at Eurofins Genomics, Karnataka, India. The adapters were designed to allow selective cleavage of the forward strands after re-synthesis of the reverse strand during sequencing. The copied reverse strands were used to sequence from the opposite end of the fragment. The sequenced raw data were processed to obtain high-quality paired-end reads using Trimmomatic v0.38 and an in-house script to remove adapters, ambiguous reads by considering parameters, *i.e.* reads that contain unknown nucleotides “N” larger than 5% was removed and low-quality reads with more than 10% quality threshold (QV) < 20 Phred score was removed (the dirty raw reads (i.e., reads with adaptors, reads with unknown nucleotides larger than 5%, low quality reads) were discarded, the average quality within the window falls below a threshold of 20 was removed, the bases of the start of a read were removed, if below a threshold quality of 20, the bases of the end of a read were removed, if below a threshold quality of 20. Based on these parameters, the clean reads were filtered from raw sequencing data and the low-quality reads containing unknown nucleotides or adaptor sequences were removed.

High quality reads from all 4 samples were pooled together and assembled into transcripts using Trinity de novo assembler (version 2.8.4) with a kmer of 25 as used in (Zhou *et al*., 2018). *De novo* assembly of the short reads (from paired-end RNA-seq library) into transcriptome was carried out with six combinations such as RC vs. RI, RC vs. SC, RC vs. SI, RI vs. SI, SC vs. RI and SC vs. SI. The assembled transcripts were then further clustered together using CD-HIT-EST-4.6 to remove the isoforms produced during assembly and resulted in sequences that can no longer be extended called as unigenes. Only those unigenes which were found to have >80% coverage at 3X read depth were considered for downstream analysis.

### Coding Sequence (CDS) Prediction and Gene annotation analysis

The coding sequences were predicted using TransDecoder-v5.3.0 from the resulted unigenes. The TransDecoder identified candidate-coding regions within unigene sequences. TransDecoder identifies likely coding sequences based on the criteria’s i.e. a minimum length open reading frame (ORF) should be found in a unigene sequence, a log-likelihood score should be > 0 and the above coding score was greatest when the ORF was scored in the 1^st^ reading frame as compared to scores in the other 5 reading frames. If a candidate ORF was found fully encapsulated by the coordinates of another candidate ORF, the longer one was reported. The sample wise CDS from the pooled set of CDS were mapped on the final set of pooled CDS using BWA (-mem) toolkit. The read count (RC) values were calculated from the resulting mapping and those CDS having 80% coverage and 3X read depth were considered for downstream analysis for each of the samples. Functional annotation of the genes was performed using DIAMOND alignment program, which is a BLAST-compatible local aligner for mapping translated DNA query sequences against a protein reference database (Buchfink *et al*., 2015). DIAMOND (BLASTX alignment mode) finds the homologous sequences for the genes against NR (non-redundant protein database) from NCBI.

### Gene Ontology Analysis

Gene ontology (GO) analyses of the CDS identified for each of the 4 leaf samples were carried out using the Blast2GO program. GO assignments were used to classify the functions of the predicted CDS. The GO mapping also provides an ontology of defined terms representing gene product properties, which are grouped into three main domains: Biological Process (BP), Molecular Function (MF) and Cellular Component (CC). GO mapping was carried out to retrieve GO terms for all the functionally annotated CDS. The GO mapping uses following criteria to retrieve GO terms for the functionally annotated CDS: BlastX result accession IDs were used to retrieve gene names or symbols, identified gene names or symbols were then searched in the species-specific entries of the gene-product tables of GO database, BlastX result accession IDs were used to retrieve UniProt IDs making use of protein information resource(PIR) which includes Protein Sequence *Database* (PSD), UniProt, SwissProt, TrEMBL, RefSeq, GenPept and Protein *Data* Bank (PDB) databases, Accession IDs were searched directly in the dbxref table of GO database.

### Functional Annotation KEGG Pathways

From this predicted CDS in the four sequence libraries, functional gene annotation analysis was performed by annotating and mapping via the Kyoto Encyclopedia of Genes and Genomes (KEGG) database. Hence, the metabolic pathway analysis of the four-leaf samples libraries was carried out using the KEGG database. The potential involvement of the predicted CDS was categorized into 23 KEGG pathways under five main categories i.e. Metabolism, Genetic information processing, Environmental information processing, Cellular processes and Organismal systems.

### Differential gene expression analysis

Differential expression analysis was performed on the CDS between control and treated samples by employing a negative binomial distribution model in DESeq package (version 1.22.1- http://www.huber.embl.de/users/anders/DESeq/). Dispersion values were estimated with the following parameters: method = blind, sharing Mode = fit-only and fit type = local. The gene expression levels were reported in transcript per million (TPM). Log2 fold change (FC) value was calculated on the read abundance using the formula: FC=Log2 (Treated/Control). The CDSs having log2foldchange value greater than zero were considered as up-regulated whereas less than zero as down-regulated at P-value threshold of 0.05.

### Validation transcriptome sequencing by RT-qPCR

We utilized the same RNA used for RNA-Seq for qPCR based validation of selected genes. The cDNA was synthesised using RevertAid First-strand cDNA synthesis kit using Oligo dT primers as per manufacturer protocol. For validation of differentially expressed genes, six differentially expressed gene associated with *P. capsici* infection were randomly selected for validation. Actin-7 like gene was used as an internal control gene for data normalization. Primers were designed using Primer Express 3.0.1 software (Table S8). The RT-qPCR reaction was performed in three biological replication for each sample using SYBR®Green Jump Start™ Taq Ready Mix™ (Sigma) on Applied Biosystems’ Step One™ Real-Time PCR System (Choudhri *et al*., 2018). The RT-qPCR analysis with a total reaction volume of 10µl containing 5µl of 2x SYBR Green JumpStart Taq ready mix, 1µl of each forward primer (10 μM) and reverse primer (10 μM), 2µl of 100ng/µL cDNA and 1µL of nuclease-free water was performed. The amplification of RT-qPCR run method was adjusted as the following conditions: the reaction system was heated to 94°C and denatured for 2 min, the main reaction (denaturation at 94°C for 15 seconds, annealing 58-60°C for 60 seconds, and template extensions at 72°C for 15 seconds) for 40cycles. The RT-qPCR product was analysed using StepOne™ Software v.2.2.2. The *C. annuum* actin-7 like gene (GenBank: GQ339766.1) was used as an internal control and expression levels between control and treated leaf samples were analysed using the 2^−ΔΔ*C*t^ method (Shen *et al*., 2015). The standard error in expression levels of three biological replications was denoted by error bars in the figures and the results were expressed as mean value ± SD. The RT-qPCR amplified product was run and visualized using 1% agarose gel electrophoresis as the method used by Gupta *et al*., 2017.

## Supporting information

https://www.ncbi.nlm.nih.gov/biosample/16251797

## Abbreviations

RC: Resistant control, RI: Resistant infected, SC: Susceptible control, Susceptible infected, TL: Transcriptome length, UL: Unigene length, UGT: Uridine 5’-diphospho glucuronosyltransferase, LTLP: lipid transfer-like protein, ERTF: ethylene-responsive transcription factor, PRP: Pathogenesis related protein, EP3: Endochitinase like protein, GRP: glycine-rich protein, CDS: coding sequence, bp: base pair, NCBI: National Center for Biotechnology Information, DEG: Differential Expression of Gene, GO: Gene Ontology, PDB: Protein Database, KEGG: Kyoto Encyclopedia of Genes and Genomes, TFDB: Transcription factor database, MYB: Myb-like DNA-binding domain, bHLH: basic helix loop helix, NAC: Pfam: Protein family, SSR: Simple sequence repeat, PE: Paired-end.

## Declarations

### Ethics approval and consent to participate

This research did not involve the use of any animal or human data or tissue.

### Consent for publication

Not applicable

## Availability of data and materials

The transcriptome sequencing data for the four-leaf samples with BioProject Id PRJNA665332, BioSample accession number RC: SAMN16251280; RI: SAMN16251798; SC: SAMN16251797; SI: SAMN16251799 were archived on SRA at the link https://www.ncbi.nlm.nih.gov/biosample/16251797

## Author contributions

Conceptualization, Data curation, Formal analysis, Investigation, Methodology, Writing an original draft, Software, Validation, T.R and V.C., Project administration, Resource, V.C, Writing-Review and editing, Supervision, Om.P.G and V.C

## Acknowledgements

We acknowledge Eurofins Genomics India Pvt. Ltd., Bengaluru, India for Illumina sequencing and Bioinformatics analysis. TR is highly thankful to the Ministry of Science and Higher Education, Ethiopia (former Ministry of Education) for sponsoring through the Fellowship Program and Department Bio and Nanotechnology, Guru Jambheshwar University of Science and Technology, Hisar, India for providing all necessary laboratory facilities.

---

**Fig S1:**
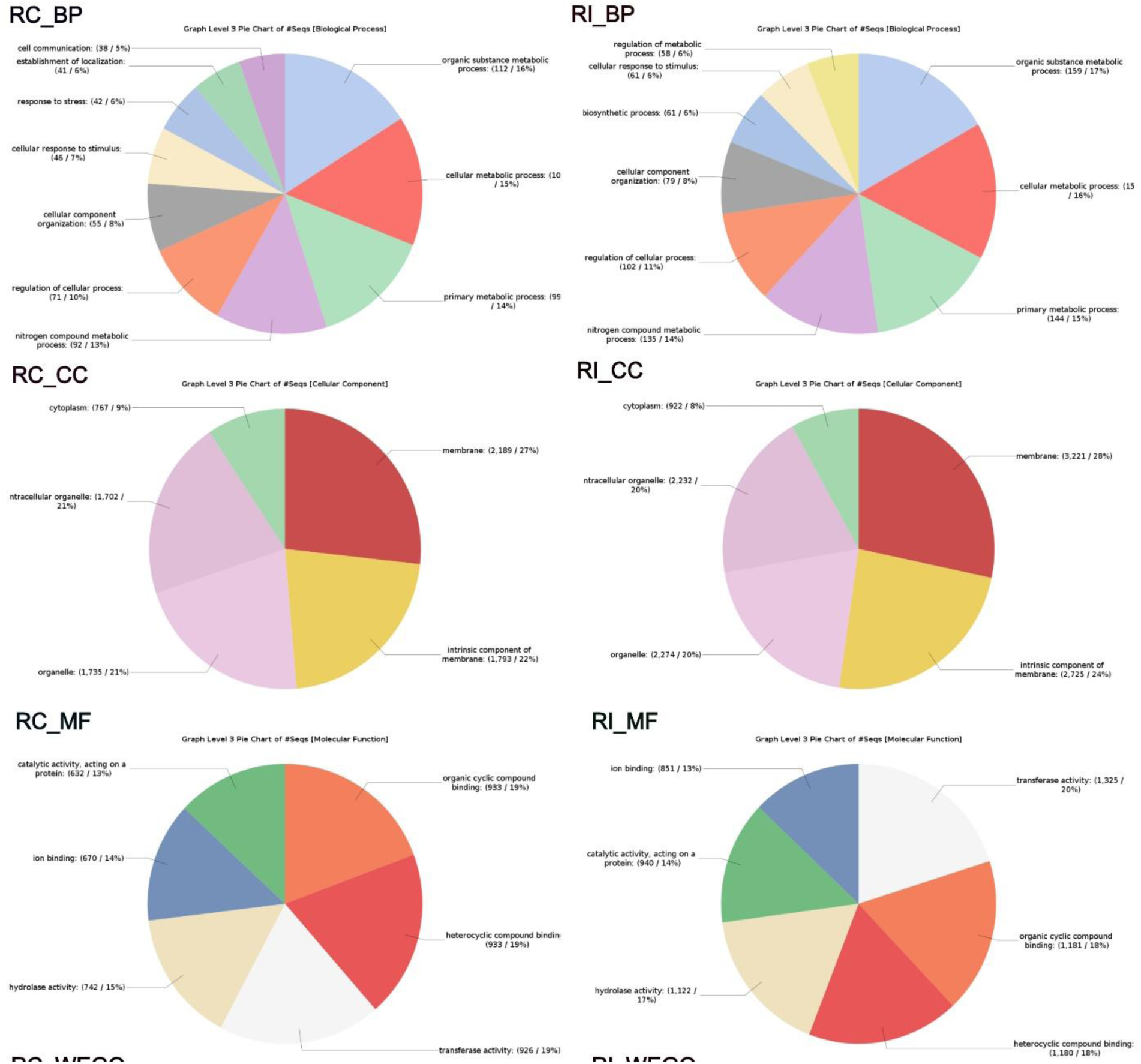
The number of predicted CDS assigned into the three GO term distribution in RC and RI leaf sample. (RI_BP, Biological processing; RI_CC, Cellular Component; RI_MF, Molecular Function).

**Fig S2:**
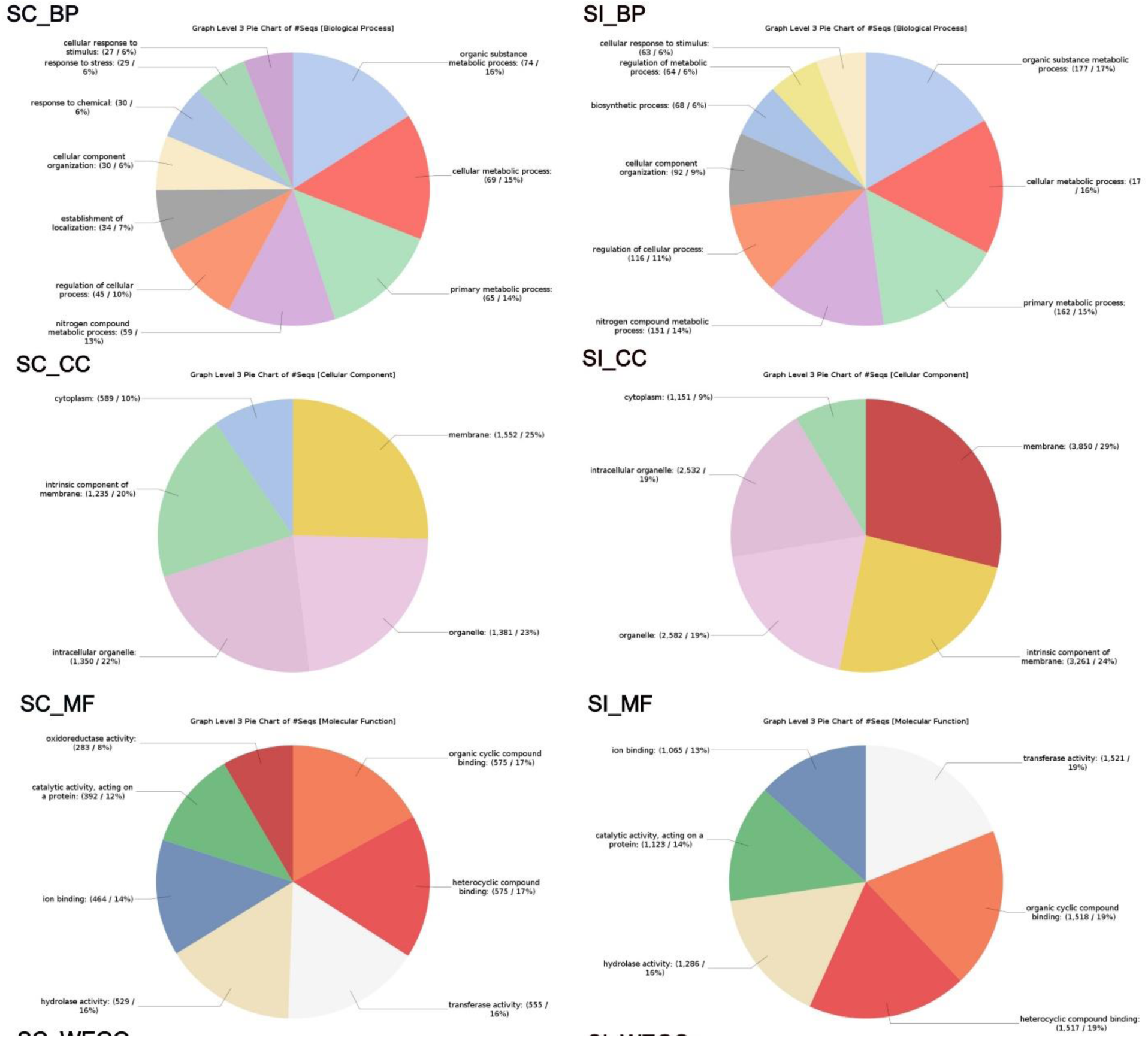
The number of predicted CDS assigned into the three GO term distribution in SC and SI leaf sample. (SC_BP, Biological processing; SC_CC, Cellular Component; SC_MF, Molecular function).

**Fig S3.**
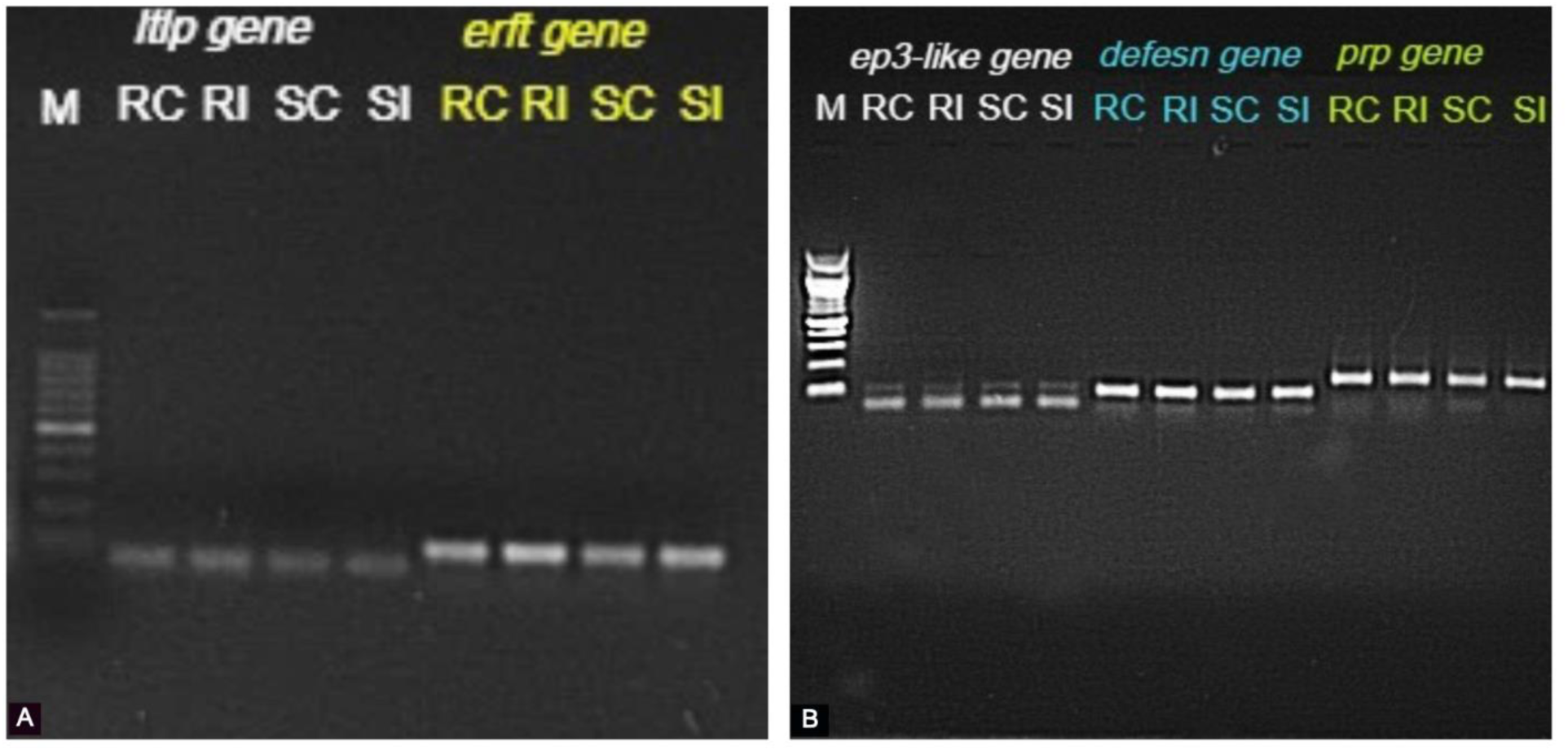
The RT-qPCR amplified product of differentially expressed genes from control and treated leaf samples, i.e. RC, RI, SC and SI run on 1% agarose gel electrophoresis; A) lipid transfer-like protein VAS, and ethylene-responsive transcription factor RAP2-7-like isoform X2genes, B) endochitinase EP3-like, defensin J1-2-like, and pathogenesis-related protein STH-2-like genes. **(Original Gel image)**. The RT-qPCR amplified product of differentially expressed genes from control and treated leaf samples, i.e. RC, RI, SC and SI run on 1% agarose gel electrophoresis; A) lipid transfer-like protein VAS, and ethylene-responsive transcription factor RAP2-7-like isoform X2 genes, B) endochitinase EP3-like, defensin J1-2-like, and pathogenesis-related protein STH-2-like genes.

**Fig S4:**
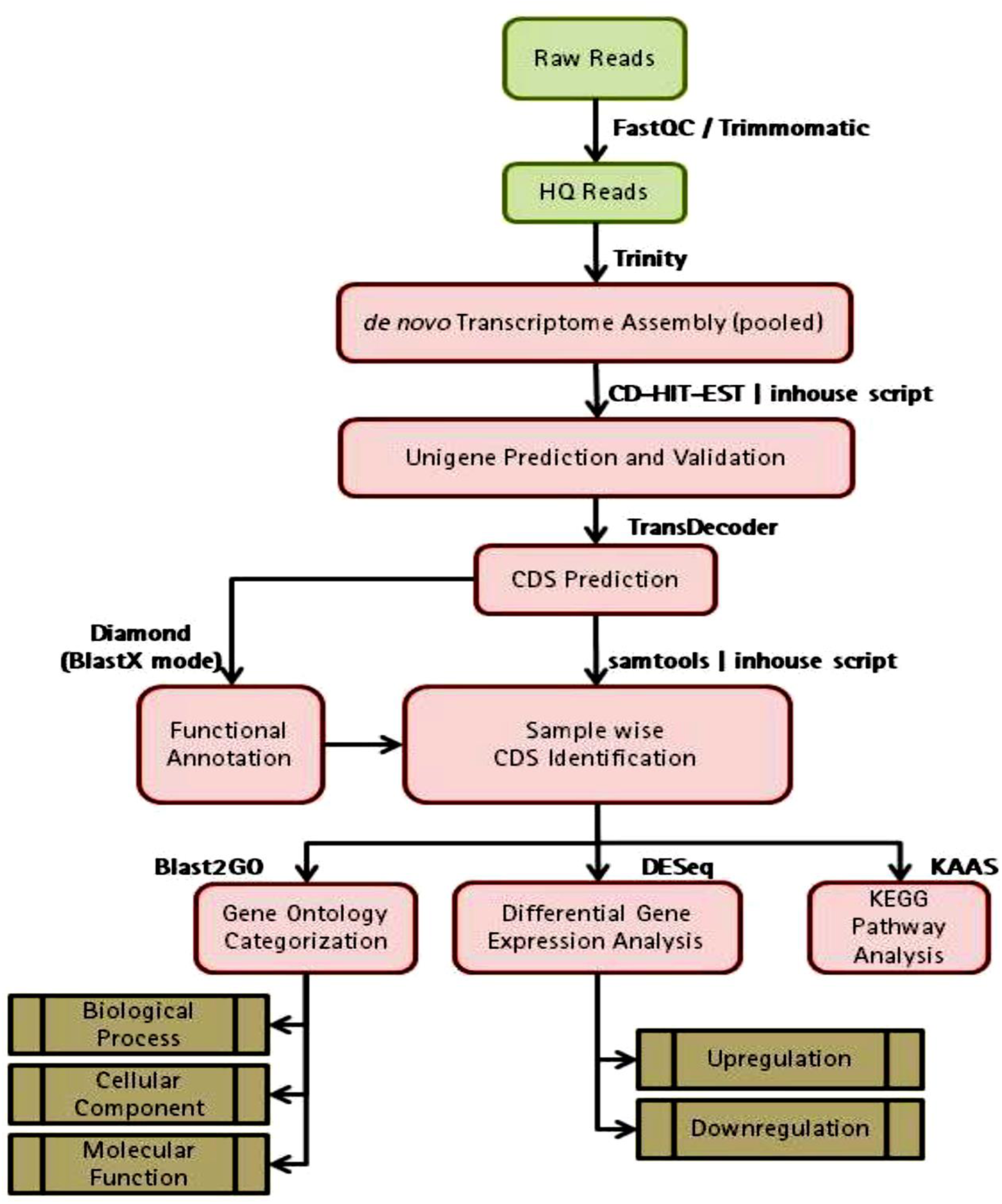
Bioinformatics Data Analysis Workflow including the steps used and bioinformatics tools undertaken during transcriptome sequencing of *Capsicum annuum* L.

**Table S1:**
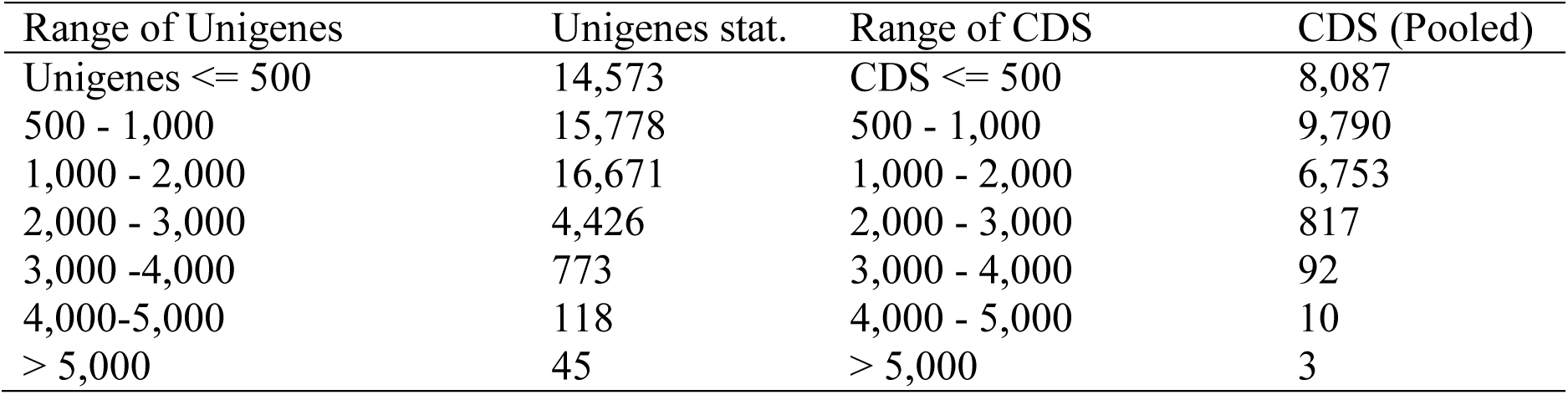
Length distribution statistics of the validated unigenes and CDS in pooled assemblies of the four-leaf sample of chili pepper (RC, RI, SC and SI).

**Table S2:**
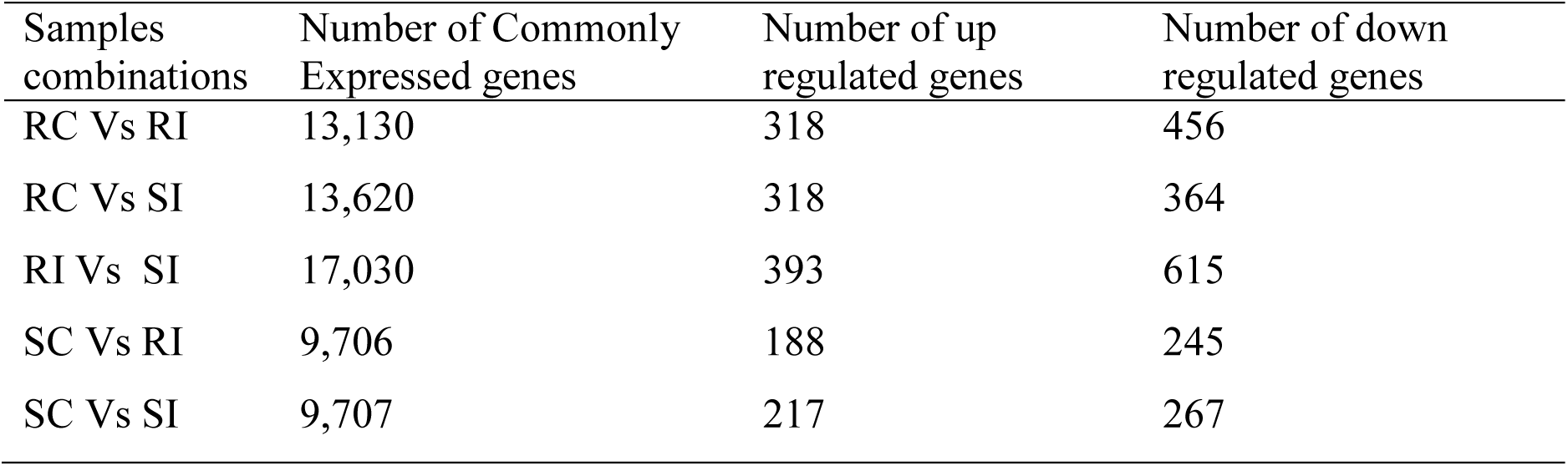
Number of commonly expressed genes, differentially expressed genes, number of up-regulated and down-regulated genes between the five combinations of control and treated leaf samples

**Table S3:**
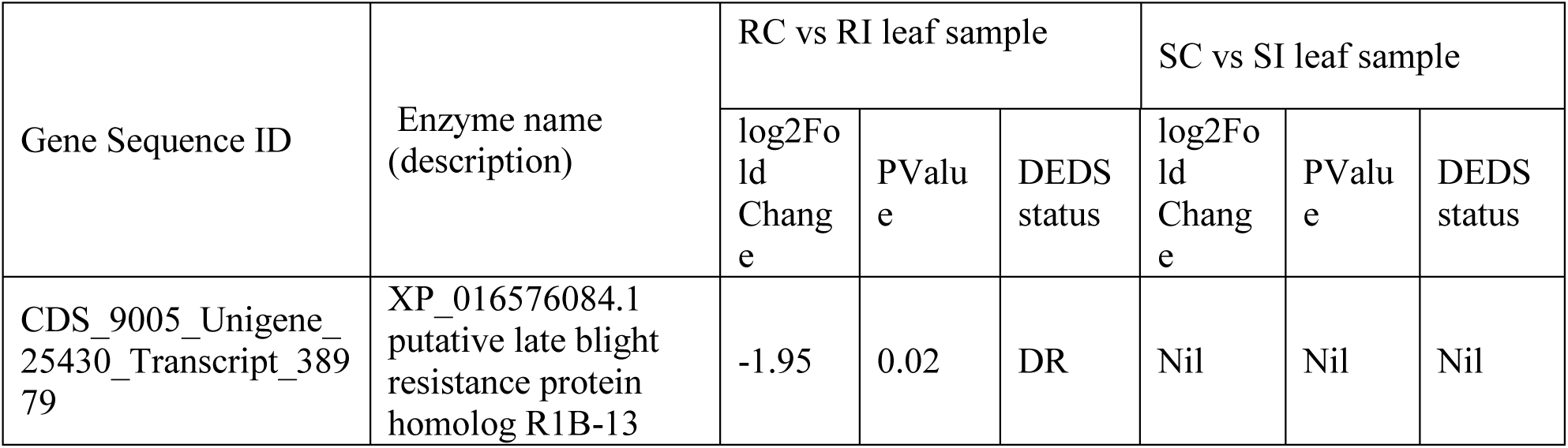
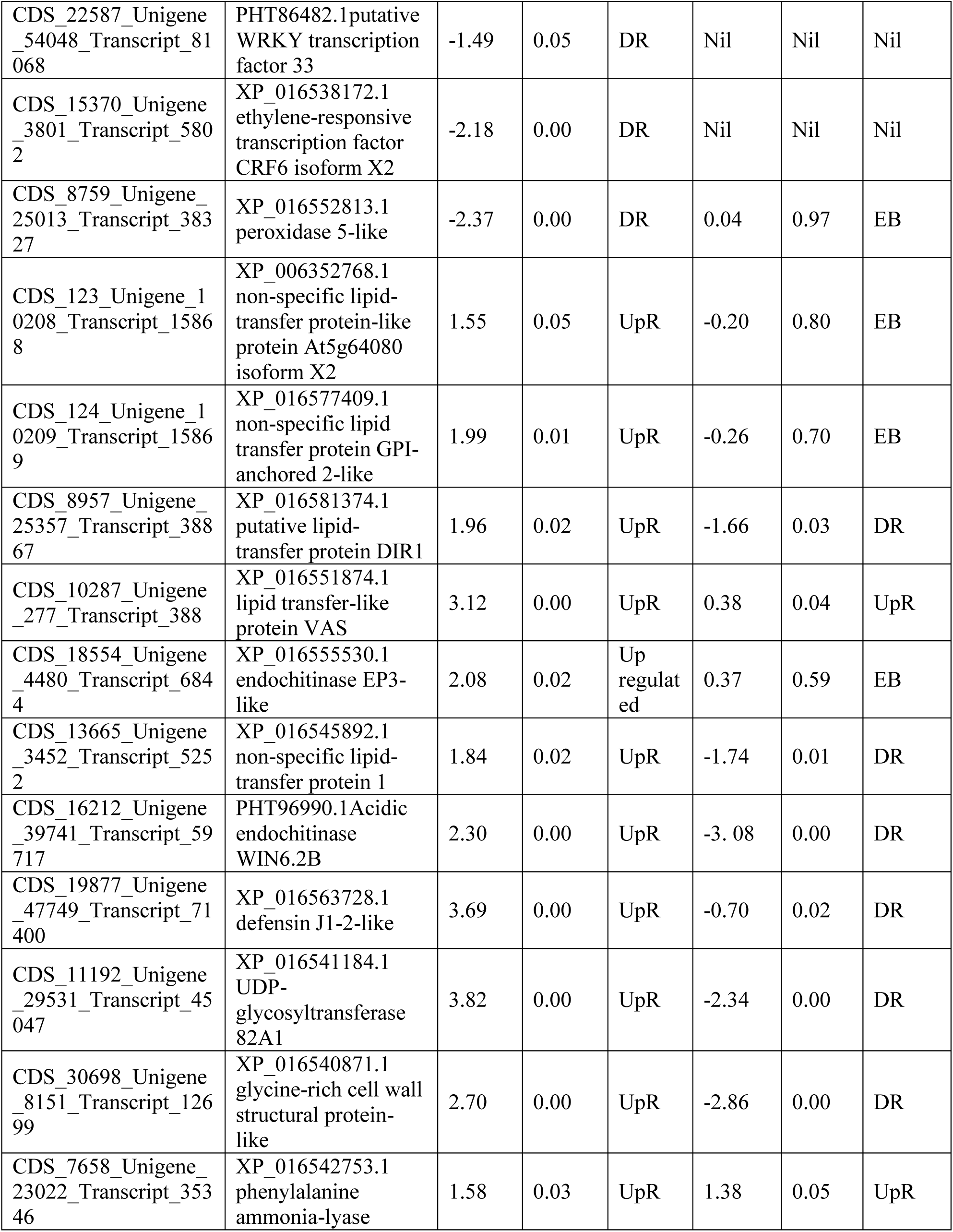
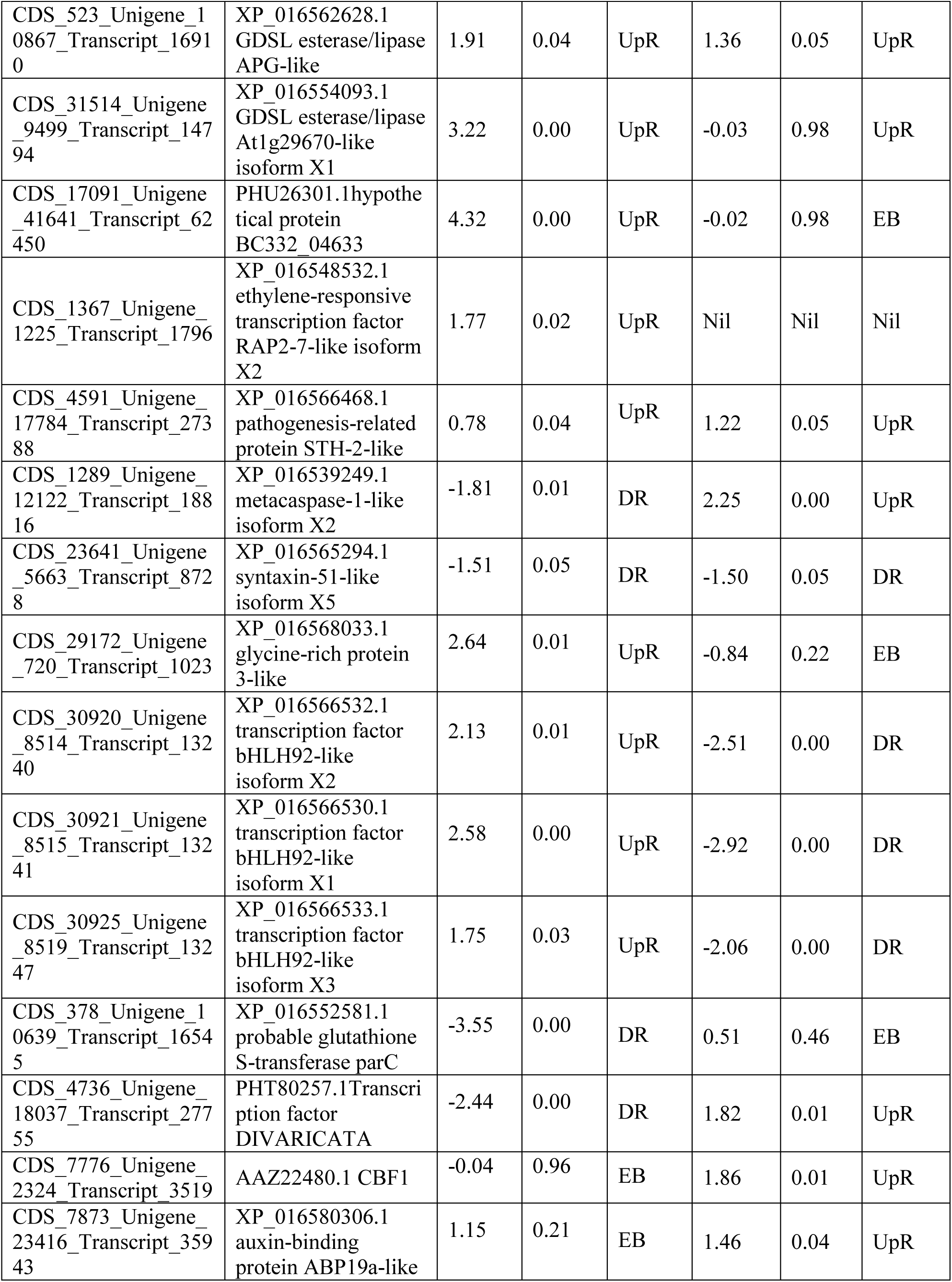
Differentially expressed genes associated with *P. capsici* infection between RC vs. RI and SC vs. SI leaf sample (DR=Down regulated, UpR=Up regulated, and EB=Expressed both).

**Table S4.**
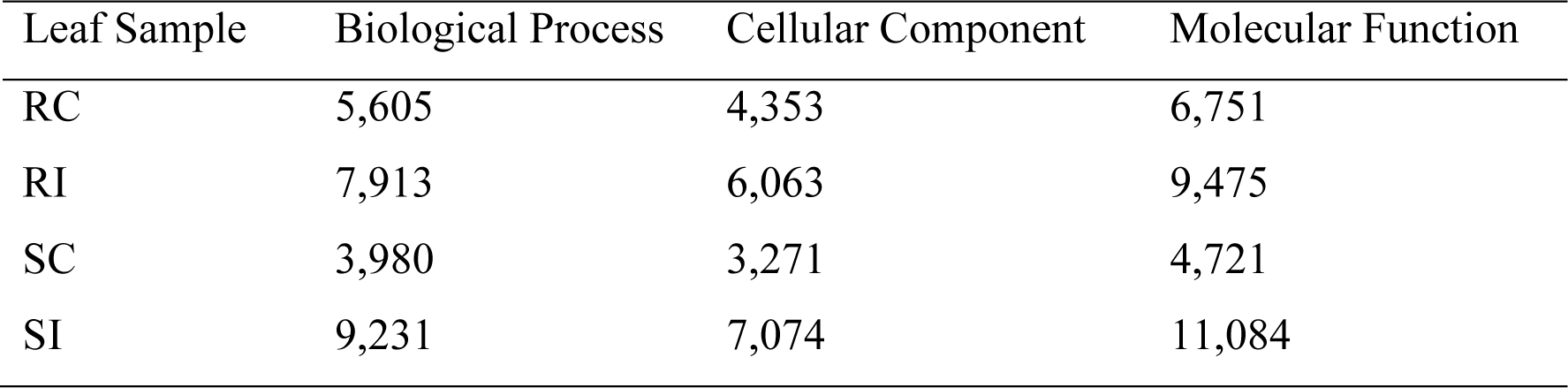
GO category distribution of CDS in RC, RI, SC and SI leaf samples.

**Table S5:**
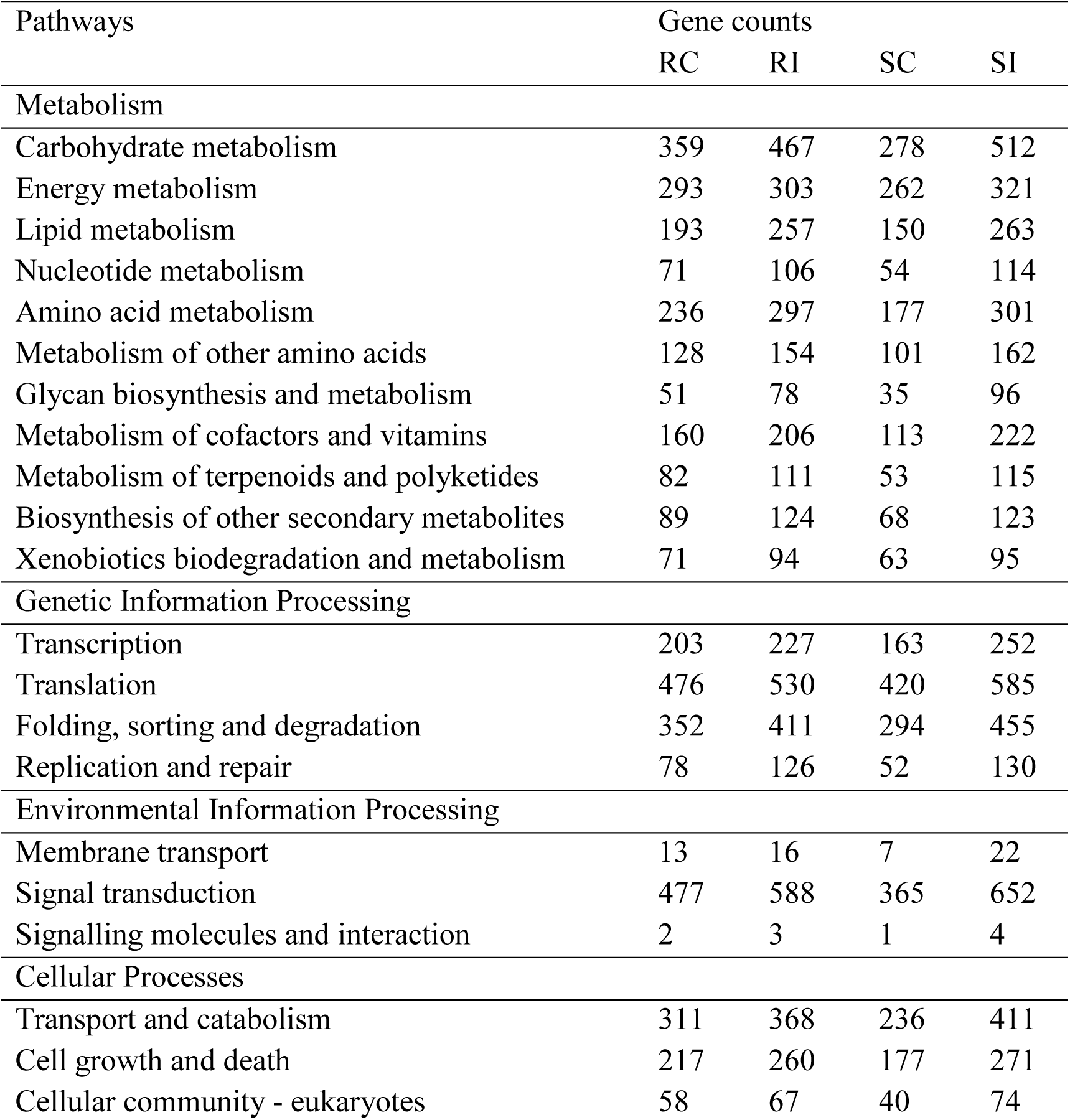
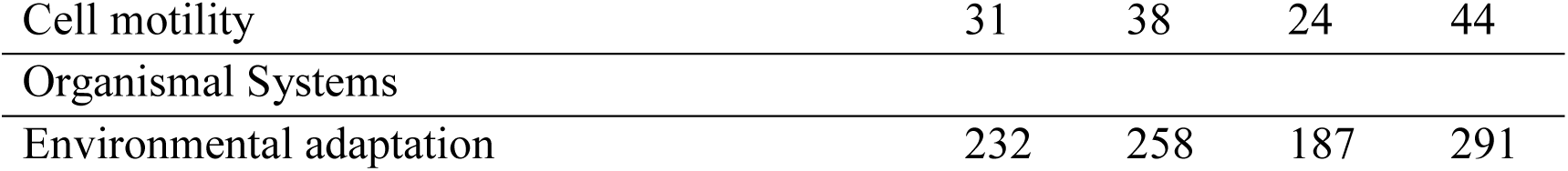
The KEGG pathways classification of CDS from the four-leaf sample of chili pepper under control and treated condition

**Table S6:**
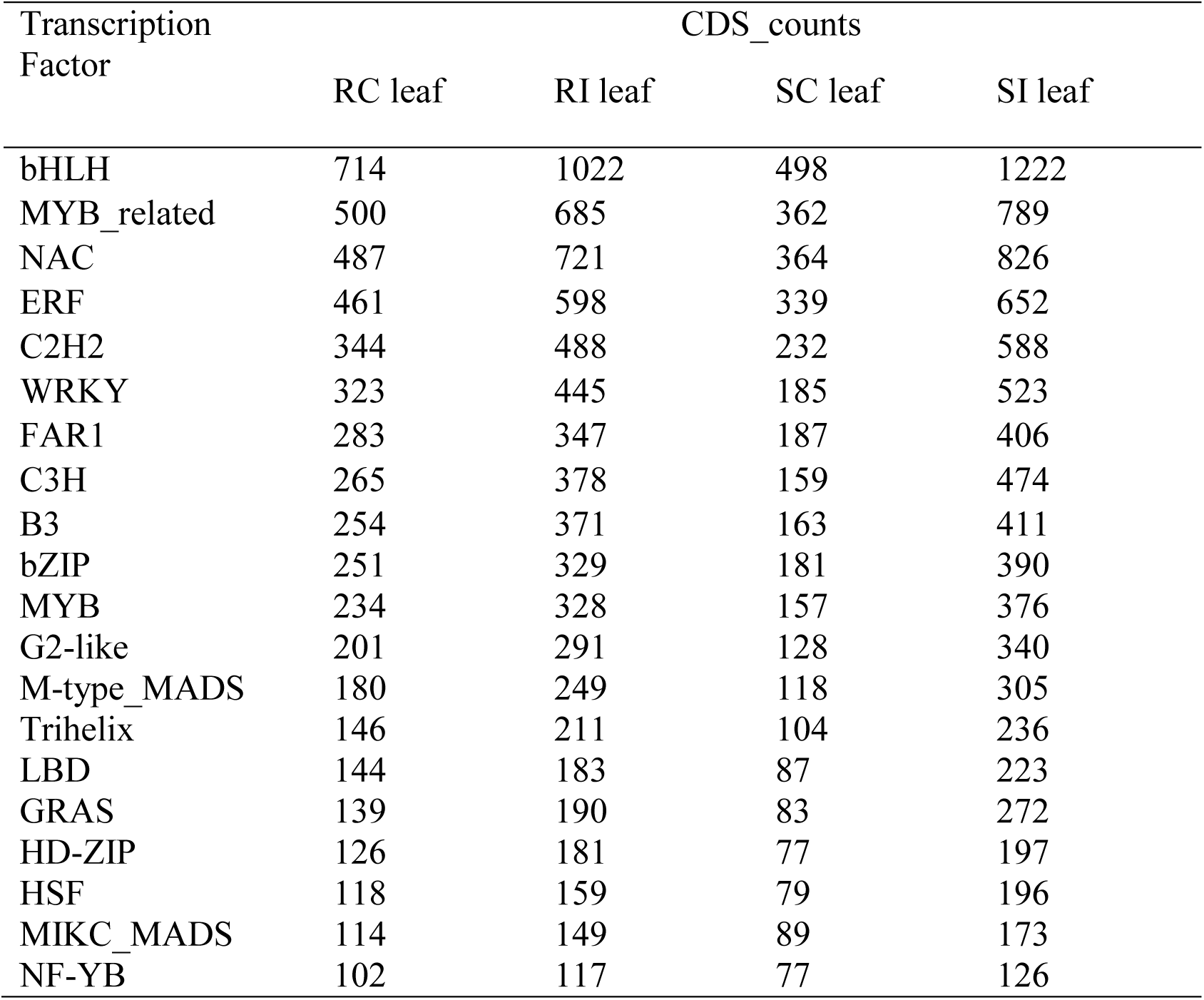
The list of 20 transcription factors enriched in the four chili pepper leaf sample under control and *P. capsici* infection condition

**Table S7:**
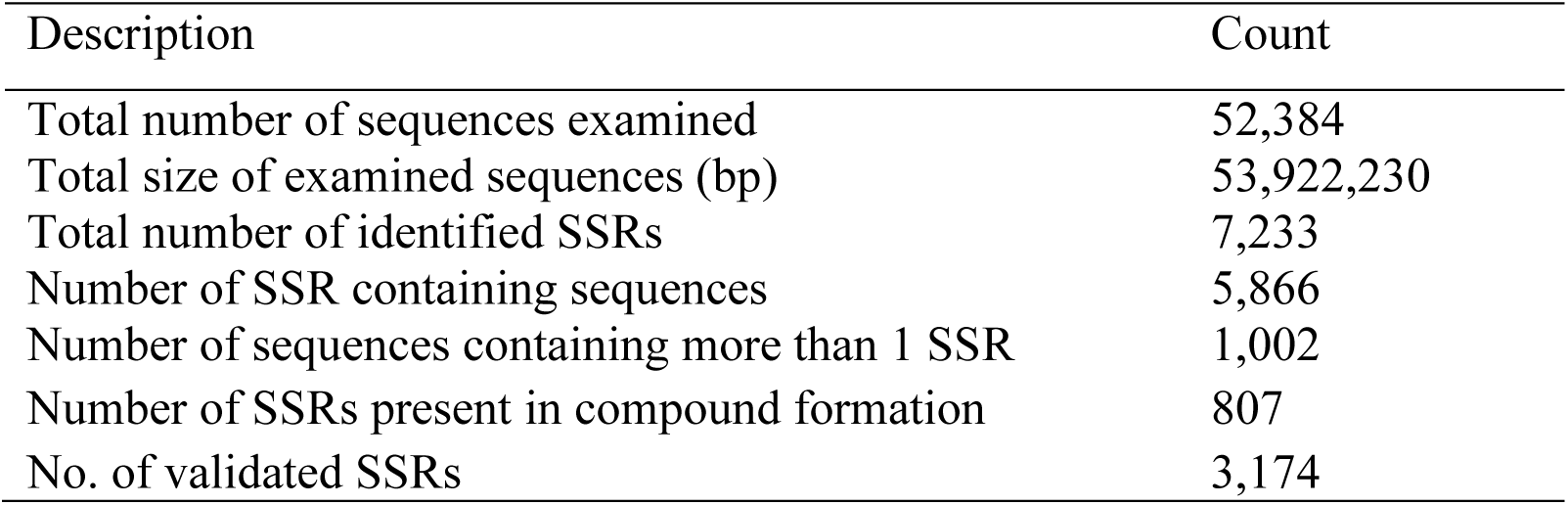
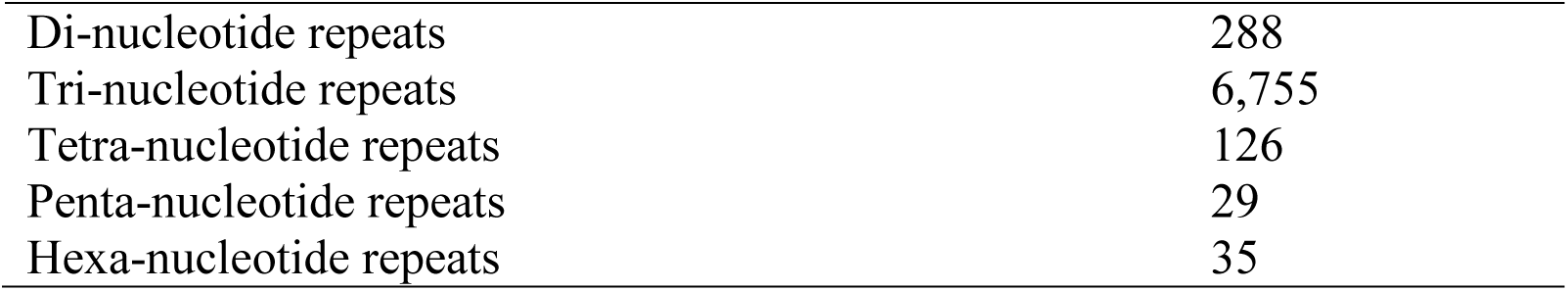
SSR Identification statistics and distribution of repeats across the four-leaf sample of chili pepper

**Table S8.**
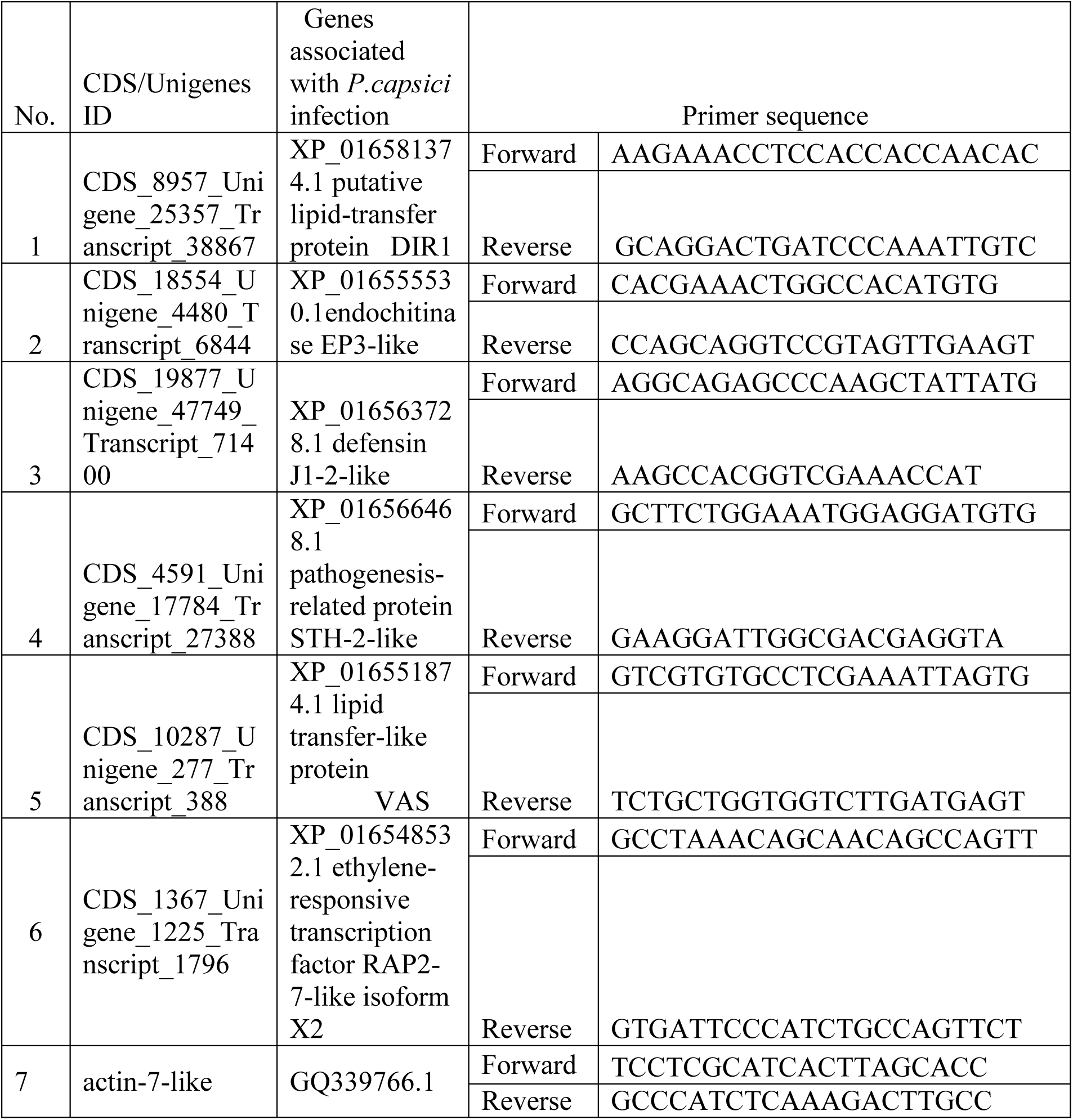
Six set of primers designed for validation of six randomly selected DEG genes associated with defence response against *P. capsici* using RT-qPCR.

