## Supplementary material for "Integrative RNA-Seq analysis of *Capsicum annuum L*.*-Phytophthora capsici L.* pathosystem reveals molecular cross-talk and activation of host defence response": https://www.ncbi.nlm.nih.gov/biosample/16251797: Supplementary tables.pdf

Table S1: Length distribution statistics of the validated unigenes and CDS in pooled assemblies of four leaf sample of chili pepper (RC, RI, SC and SI).

| Range of Unigenes | Unigenes stat. | Range of CDS | CDS (Pooled) |
|---|---|---|---|
| Unigenes <= 500 | 14,573 | CDS <= 500 | 8,087 |
| 500 - 1,000 | 15,778 | 500 - 1,000 | 9,790 |
| 1,000 - 2,000 | 16,671 | 1,000 - 2,000 | 6,753 |
| 2,000 - 3,000 | 4,426 | 2,000 - 3,000 | 817 |
| 3,000 -4,000 | 773 | 3,000 - 4,000 | 92 |
| 4,000-5,000 | 118 | 4,000 - 5,000 | 10 |
| > 5,000 | 45 | > 5,000 | 3 |

Table S2: Number of commonly expressed genes, differentially expressed genes, number of up regulated and down regulated genes between the five combinations of control and treated leaf samples

| Samples combinations | Number of Commonly Expressed genes | Number of up regulated genes | Number of down regulated genes |
|---|---|---|---|
| RC Vs RI | 13,130 | 318 | 456 |
| RC Vs SI | 13,620 | 318 | 364 |
| RI Vs SI | 17,030 | 393 | 615 |
| SC Vs RI | 9,706 | 188 | 245 |
| SC Vs SI | 9,707 | 217 | 267 |

Table S3. Differentially expressed genes associated to *P. capsici* infection between RC vs RI and SC vs SI leaf sample (DR=Down regulated, UpR=Up regulated, and EB=Expressed both).

| Gene Sequence ID | Enzyme name (description) | RC vs RI leaf sample | | | SC vs SI leaf sample | | |
|---|---|---|---|---|---|---|---|
| | | log2Fold Change | PValue | DEDS status | log2Fold Change | PValue | DEDS status |
| CDS_9005_Unigene_25430_Transcript_38979 | XP_016576084.1 putative late blight resistance protein homolog R1B-13 | -1.95 | 0.02 | DR | Nil | Nil | Nil |
| CDS_22587_Unigene_54048_Transcript_81068 | PHT86482.1putative WRKY transcription factor 33 | -1.49 | 0.05 | DR | Nil | Nil | Nil |
| CDS_15370_Unigene_3801_Transcript_5802 | XP_016538172.1 ethylene-responsive transcription factor CRF6 isoform X2 | -2.18 | 0.00 | DR | Nil | Nil | Nil |

| | | | | | | | |
|---|---|---|---|---|---|---|---|
| CDS_8759_Unigene_25013_Transcript_38327 | XP_016552813.1 peroxidase 5-like | -2.37 | 0.00 | DR | 0.04 | 0.97 | EB |
| CDS_123_Unigene_10208_Transcript_15868 | XP_006352768.1 non-specific lipid-transfer protein-like protein At5g64080 isoform X2 | 1.55 | 0.05 | UpR | -0.20 | 0.80 | EB |
| CDS_124_Unigene_10209_Transcript_15869 | XP_016577409.1 non-specific lipid transfer protein GPI-anchored 2-like | 1.99 | 0.01 | UpR | -0.26 | 0.70 | EB |
| CDS_8957_Unigene_25357_Transcript_38867 | XP_016581374.1 putative lipid-transfer protein DIR1 | 1.96 | 0.02 | UpR | -1.66 | 0.03 | DR |
| CDS_10287_Unigene_277_Transcript_388 | XP_016551874.1 lipid transfer-like protein VAS | 3.12 | 0.00 | UpR | 0.38 | 0.04 | UpR |
| CDS_18554_Unigene_4480_Transcript_6844 | XP_016555530.1 endochitinase EP3-like | 2.08 | 0.02 | Up regulated | 0.37 | 0.59 | EB |
| CDS_13665_Unigene_3452_Transcript_5252 | XP_016545892.1 non-specific lipid-transfer protein 1 | 1.84 | 0.02 | UpR | -1.74 | 0.01 | DR |
| CDS_16212_Unigene_39741_Transcript_59717 | PHT96990.1Acidic endochitinase WIN6.2B | 2.30 | 0.00 | UpR | -3. 08 | 0.00 | DR |
| CDS_19877_Unigene_47749_Transcript_71400 | XP_016563728.1 defensin J1-2-like | 3.69 | 0.00 | UpR | -0.70 | 0.02 | DR |
| CDS_11192_Unigene_29531_Transcript_45047 | XP_016541184.1 UDP-glycosyltransferase 82A1 | 3.82 | 0.00 | UpR | -2.34 | 0.00 | DR |
| CDS_30698_Unigene_8151_Transcript_12699 | XP_016540871.1 glycine-rich cell wall structural protein-like | 2.70 | 0.00 | UpR | -2.86 | 0.00 | DR |
| CDS_7658_Unigene_23022_Transcript_35346 | XP_016542753.1 phenylalanine ammonia-lyase | 1.58 | 0.03 | UpR | 1.38 | 0.05 | UpR |
| CDS_523_Unigene_10867_Transcript_16910 | XP_016562628.1 GDSL esterase/lipase APG-like | 1.91 | 0.04 | UpR | 1.36 | 0.05 | UpR |
| CDS_31514_Unigene_9499_Transcript_14794 | XP_016554093.1 GDSL esterase/lipase At1g29670-like isoform X1 | 3.22 | 0.00 | UpR | -0.03 | 0.98 | UpR |
| CDS_17091_Unigene_41641_Transcript_62450 | PHU26301.1hypothetical protein BC332_04633 | 4.32 | 0.00 | UpR | -0.02 | 0.98 | EB |
| CDS_1367_Unigene_1225_Transcript_1796 | XP_016548532.1 ethylene-responsive transcription factor | 1.77 | 0.02 | UpR | Nil | Nil | Nil |

| | | | | | | | |
|---|---|---|---|---|---|---|---|
| | RAP2-7-like isoform X2 | | | | | | |
| CDS_4591_Unigene_17784_Transcript_27388 | XP_016566468.1 pathogenesis-related protein STH-2-like | 0.78 | 0.04 | UpR | 1.22 | 0.05 | UpR |
| CDS_1289_Unigene_12122_Transcript_18816 | XP_016539249.1 metacaspase-1-like isoform X2 | -1.81 | 0.01 | DR | 2.25 | 0.00 | UpR |
| CDS_23641_Unigene_5663_Transcript_8728 | XP_016565294.1 syntaxin-51-like isoform X5 | -1.51 | 0.05 | DR | -1.50 | 0.05 | DR |
| CDS_29172_Unigene_720_Transcript_1023 | XP_016568033.1 glycine-rich protein 3-like | 2.64 | 0.01 | UpR | -0.84 | 0.22 | EB |
| CDS_30920_Unigene_8514_Transcript_13240 | XP_016566532.1 transcription factor bHLH92-like isoform X2 | 2.13 | 0.01 | UpR | -2.51 | 0.00 | DR |
| CDS_30921_Unigene_8515_Transcript_13241 | XP_016566530.1 transcription factor bHLH92-like isoform X1 | 2.58 | 0.00 | UpR | -2.92 | 0.00 | DR |
| CDS_30925_Unigene_8519_Transcript_13247 | XP_016566533.1 transcription factor bHLH92-like isoform X3 | 1.75 | 0.03 | UpR | -2.06 | 0.00 | DR |
| CDS_378_Unigene_10639_Transcript_16545 | XP_016552581.1 probable glutathione S-transferase parC | -3.55 | 0.00 | DR | 0.51 | 0.46 | EB |
| CDS_4736_Unigene_18037_Transcript_27755 | PHT80257.1Transcription factor DIVARICATA | -2.44 | 0.00 | DR | 1.82 | 0.01 | UpR |
| CDS_7776_Unigene_2324_Transcript_3519 | AAZ22480.1 CBF1 | -0.04 | 0.96 | EB | 1.86 | 0.01 | UpR |
| CDS_7873_Unigene_23416_Transcript_35943 | XP_016580306.1 auxin-binding protein ABP19a-like | 1.15 | 0.21 | EB | 1.46 | 0.04 | UpR |

Table S4: GO category distribution of CDS in RC, RI, SC and SI leaf samples

| Leaf Sample | Biological Process | Cellular Component | Molecular Function |
|---|---|---|---|
| RC | 5,605 | 4,353 | 6,751 |
| RI | 7,913 | 6,063 | 9,475 |
| SC | 3,980 | 3,271 | 4,721 |
| SI | 9,231 | 7,074 | 11,084 |

Table S5: The KEGG pathways classification of CDS from four-leaf sample of chili pepper under control and treated condition

| Pathways | Gene counts | | | |
|---|---|---|---|---|
| | RC | RI | SC | SI |
| Metabolism | | | | |
| Carbohydrate metabolism | 359 | 467 | 278 | 512 |
| Energy metabolism | 293 | 303 | 262 | 321 |
| Lipid metabolism | 193 | 257 | 150 | 263 |
| Nucleotide metabolism | 71 | 106 | 54 | 114 |
| Amino acid metabolism | 236 | 297 | 177 | 301 |
| Metabolism of other amino acids | 128 | 154 | 101 | 162 |
| Glycan biosynthesis and metabolism | 51 | 78 | 35 | 96 |
| Metabolism of cofactors and vitamins | 160 | 206 | 113 | 222 |
| Metabolism of terpenoids and polyketides | 82 | 111 | 53 | 115 |
| Biosynthesis of other secondary metabolites | 89 | 124 | 68 | 123 |
| Xenobiotics biodegradation and metabolism | 71 | 94 | 63 | 95 |
| Genetic Information Processing | | | | |
| Transcription | 203 | 227 | 163 | 252 |
| Translation | 476 | 530 | 420 | 585 |
| Folding, sorting and degradation | 352 | 411 | 294 | 455 |
| Replication and repair | 78 | 126 | 52 | 130 |
| Environmental Information Processing | | | | |
| Membrane transport | 13 | 16 | 7 | 22 |
| Signal transduction | 477 | 588 | 365 | 652 |
| Signalling molecules and interaction | 2 | 3 | 1 | 4 |
| Cellular Processes | | | | |
| Transport and catabolism | 311 | 368 | 236 | 411 |
| Cell growth and death | 217 | 260 | 177 | 271 |
| Cellular community - eukaryotes | 58 | 67 | 40 | 74 |
| Cell motility | 31 | 38 | 24 | 44 |
| Organismal Systems | | | | |
| Environmental adaptation | 232 | 258 | 187 | 291 |

Table S6: The list of 20 transcription factors enriched in the four chili pepper leaf sample under control and *P. capsici* infection condition

| Transcription Factor | CDS_counts | | | |
|---|---|---|---|---|
| | RC leaf | RI leaf | SC leaf | SI leaf |
| bHLH | 714 | 1022 | 498 | 1222 |
| MYB_related | 500 | 685 | 362 | 789 |
| NAC | 487 | 721 | 364 | 826 |

| | | | | |
|---|---|---|---|---|
| ERF | 461 | 598 | 339 | 652 |
| C2H2 | 344 | 488 | 232 | 588 |
| WRKY | 323 | 445 | 185 | 523 |
| FAR1 | 283 | 347 | 187 | 406 |
| C3H | 265 | 378 | 159 | 474 |
| B3 | 254 | 371 | 163 | 411 |
| bZIP | 251 | 329 | 181 | 390 |
| MYB | 234 | 328 | 157 | 376 |
| G2-like | 201 | 291 | 128 | 340 |
| M-type_MADS | 180 | 249 | 118 | 305 |
| Trihelix | 146 | 211 | 104 | 236 |
| LBD | 144 | 183 | 87 | 223 |
| GRAS | 139 | 190 | 83 | 272 |
| HD-ZIP | 126 | 181 | 77 | 197 |
| HSF | 118 | 159 | 79 | 196 |
| MIKC_MADS | 114 | 149 | 89 | 173 |
| NF-YB | 102 | 117 | 77 | 126 |

Table S7. SSR Identification statistics and distribution of repeats across the four leaf sample of chili pepper.

| Description | Count |
|---|---|
| Total number of sequences examined | 52,384 |
| Total size of examined sequences (bp) | 53,922,230 |
| Total number of identified SSRs | 7,233 |
| Number of SSR containing sequences | 5,866 |
| Number of sequences containing more than 1 SSR | 1,002 |
| Number of SSRs present in compound formation | 807 |
| No. of validated SSRs | 3,174 |
| Di-nucleotide repeats | 288 |
| Tri-nucleotide repeats | 6,755 |
| Tetra-nucleotide repeats | 126 |
| Penta-nucleotide repeats | 29 |
| Hexa-nucleotide repeats | 35 |

Table S8. Six set of primers designed for validation of six randomly selected DEG genes associated with defense response against *P. capsici* using RT- qPCR.

| No. | CDS/Unigenes ID | Genes associated with *P.capsici* infection | Primer sequence | |
|---|---|---|---|---|
| 1 | CDS_8957_Unigene_25357_Transcript_38867 | XP_016581374.1 putative lipid-transfer protein DIR1 | Forward | AAGAAACCTCCACCACCAACAC |
| | | | Reverse | GCAGGACTGATCCCAAATTGTC |
| 2 | CDS_18554_Unigene_4480_Transcript_6844 | XP_016555530.1endochitinase EP3-like | Forward | CACGAAACTGGCCACATGTG |
| | | | Reverse | CCAGCAGGTCCGTAGTTGAAGT |
| 3 | CDS_19877_Unigene_47749_Transcript_71400 | XP_016563728.1 defensin J1-2-like | Forward | AGGCAGAGCCCAAGCTATTATG |
| | | | Reverse | AAGCCACGGTCGAAACCAT |
| 4 | CDS_4591_Unigene_17784_Transcript_27388 | XP_016566468.1 pathogenesis-related protein STH-2-like | Forward | GCTTCTGGAAATGGAGGATGTG |
| | | | Reverse | GAAGGATTGGCGACGAGGTA |
| 5 | CDS_10287_Unigene_277_Transcript_388 | XP_016551874.1 lipid transfer-like protein VAS | Forward | GTCGTGTGCCTCGAAATTAGTG |
| | | | Reverse | TCTGCTGGTGGTCTTGATGAGT |
| 6 | CDS_1367_Unigene_1225_Transcript_1796 | XP_016548532.1 ethylene-responsive transcription factor RAP2-7-like isoform X2 | Forward | GCCTAAACAGCAACAGCCAGTT |
| | | | Reverse | GTGATTCCCATCTGCCAGTTCT |
| 7 | actin-7-like | GQ339766.1 | Forward | TCCTCGCATCACTTAGCACC |
| | | | Reverse | GCCCATCTCAAAGACTTGCC |
