## Supplementary material for "Integrative RNA-Seq analysis of *Capsicum annuum L*.*-Phytophthora capsici L.* pathosystem reveals molecular cross-talk and activation of host defence response": https://www.ncbi.nlm.nih.gov/biosample/16251797: Transcriptom seq.Tables.pdf

**Table 1**. Statistical analysis of RNA-Seq libraries constructed from two contrasting landraces *i.e.* GojamMecha_9086 (resistant) and Dabat_80045 (susceptible) chilli pepper leaf samples exposed to *P. capsici* infection.

| **Parameters** | **GojamMecha_9086 (Resistant)** | | **Dabat_80045 (Susceptible)** | |
|---|---|---|---|---|
| | **Control** | **Infected** | **Control** | **Infected** |
| High quality reads | 19,480,516 | 25,078,845 | 22,055,840 | 22,708,051 |
| Number of bases | 2,930,503,524 | 3,773,086,364 | 3,318,642,109 | 3,415,521,810 |
| Total data (in Gb) | 2.93 | 3.77 | 3.31 | 3.41 |
| #CDS | 14,509 | 19,250 | 10,100 | 22,719 |
| Total CDS length(in bp) | 11,088,237 | 15,877,689 | 7,045,395 | 19,950,423 |
| Maximum CDS length | 13,542 | 4,878 | 13,542 | 13542 |
| Minimum CDS length | 276 | 258 | 264 | 264 |
| Mean CDS length | 764 | 825 | 698 | 878 |

| **Pooled assembly statistics** | | | | | |
|---|---|---|---|---|---|
| **Transcript description** | **Transcript stat** | **Unigene** | **Unigene stat** | **CDS** | **CDS stat** |
| Total transcripts | 1,18,879 | Total unigene | 52,384 | Total CDS | 25,552 |
| Total transcript length (bp) | 96,675,995 | Total unigene length (bp) | 53,922,230 | Total CDS length (bp) | 21,689,736 |
| N50 | 1,277 | N50 | 1,403 | Maximum CDS length (bp) | 13,542 |
| Maximum transcript length (bp) | 14,668 | Maximum unigene length (bp) | 14,668 | Minimum CDS length (bp) | 13,542 |
| Minimum transcript length (bp) | 180 | Minimum unigene length (bp) | 180 | Mean CDS length (bp) | 848.84 |
| Mean Transcript Length (bp) | 813.23 | Mean unigene length (bp) | 1029.36 | | |
